# Rapid and dynamic alternative splicing impacts the Arabidopsis cold response transcriptome

**DOI:** 10.1101/251876

**Authors:** Cristiane P. G. Calixto, Wenbin Guo, Allan B. James, Nikoleta A. Tzioutziou, Juan Carlos Entizne, Paige E. Panter, Heather Knight, Hugh G. Nimmo, Runxuan Zhang, John W. S. Brown

**Affiliations:** Plant Sciences Division, School of Life Sciences, University of Dundee, Invergowrie, Dundee, Scotland, UK.; Information and Computational Sciences, The James Hutton Institute, Invergowrie, Dundee, Scotland, UK.; Institute of Molecular, Cell and Systems Biology, College of Medical, Veterinary and Life Sciences, University of Glasgow, Glasgow, Scotland.; Cell and Molecular Sciences, The James Hutton Institute, Invergowrie, Dundee, Scotland, UK.; Department of Biosciences, Durham University, Durham DH1 3LE, UK.

**Author notes:** Equal contributors (Cristiane P. G. Calixto, Wenbin Guo).

**Keywords:** *Arabidopsis thaliana*, Differential alternative splicing, Ultra-deep RNA-seq, Time-series RNA-seq, Diel time-series, Cold response, Cold acclimation, Splicing factors, Transcription factors, Dynamics of alternative splicing.

## Abstract

**Background:** Plants have adapted to tolerate and survive constantly changing environmental conditions by re-programming gene expression. The scale of the contribution of alternative splicing (AS) to stress responses has been underestimated due to limitations in RNA-seq analysis programs and poor representation of AS transcripts in plant databases. Significantly, the dynamics of the AS response have not been investigated but this is now possible with accurate transcript quantification programs and AtRTD2, a new, comprehensive transcriptome for Arabidopsis.

**Results:** Using ultra-deep RNA-sequencing of a time-course of *Arabidopsis thaliana* plants exposed to cold treatment, we identified 8,949 genes with altered expression of which 2,442 showed significant differential alternative splicing (DAS) and 1,647 genes were regulated only at the level of AS (DAS-only). The high temporal resolution demonstrated the rapid induction of both transcription and AS resulting in coincident waves of differential expression (transcription) and differential alternative splicing in the first 6-9 hours of cold. The differentially expressed and DAS gene sets were largely non-overlapping, each comprising thousands of genes. The dynamic analysis of AS identified genes with rapid and sensitive AS within 3 h of transfer to the cold (early AS genes), which were enriched for splicing and transcription factors. A detailed investigation of the novel cold-response DAS-only gene, *U2B”-LIKE*, suggested that it regulates AS and is required for tolerance to freezing.

**Conclusions:** Our data indicate that transcription and AS are the major regulators of transcriptome reprogramming that together govern the physiological and survival responses of plants to low temperature.

## Background

Plants adapt to and survive adverse environmental conditions by reprogramming their transcriptome and proteome. Low temperatures negatively affect growth and development but, in general, plants from temperate climatic regions can tolerate chilling temperatures (0-15°C), and can increase their freezing tolerance by prior exposure to low, non-freezing temperatures through a process called cold acclimation. Cold acclimation involves multiple physiological, biochemical and molecular changes to reduce growth, modify membrane properties, and produce the proteins and metabolites required to protect cell integrity throughout the period of freezing exposure [1-3]. In *Arabidopsis thaliana*, these cold-responsive changes reflect complex reprogramming of gene expression involving chromatin modification, transcription, post-transcriptional processing, post-translational modification and protein turnover [1-5]. Previous genome-wide microarray and RNA-seq studies have mostly focussed on the cold-induced transcriptional response and thousands of differentially expressed genes have been reported. The best-studied is the signalling cascade involving the C-REPEAT-BINDING FACTOR/DEHYDRATION RESPONSE ELEMENT-BINDING PROTEIN (CBF/DREB) transcription factors (CBF1-3), which recognise the C-repeat(CRT)/dehydration responsive element (DRE) motifs in the promoters of their target genes [1, 3, 6, 7]. Increased expression of these genes during cold acclimation increases freezing tolerance, which can also be achieved by ectopic expression of the CBFs in the absence of cold acclimation [1-3, 6]. Besides the CBF regulon, other transcriptional pathways are required for the activation of cascades of gene expression which together drive the cold response, acclimation and plant survival [8]. Plants also adapt to daily and seasonal changes in temperature, and the circadian clock provides a mechanism by which they anticipate the predictable diel cycles of temperature and light/dark to optimise gene expression and consequent physiology [9]. The circadian clock also regulates cold-responsive gene expression through a process called gating where the magnitude of changes in gene expression depends on the time of day that the stimulus is applied [1, 9].

Alternative splicing (AS) has been linked to stress responses [10-17] but little is known about its global contribution or dynamics. AS is a regulated process that produces different mRNA transcripts from precursor messenger RNAs (pre-mRNAs) of a single gene [18, 19]. The selection of splice sites is determined by sequence elements in the pre-mRNA that are recognised by trans-acting splicing factors (SFs) which recruit the spliceosome for intron removal. The concentration, localisation and activity of these factors in different cells and conditions defines splice site choice to generate different alternatively spliced isoforms and splicing patterns [18, 19]. AS regulates the level of expression of genes by generating non-productive transcript isoforms which are targeted for nonsense-mediated decay (NMD), or impacts the proteome by generating protein variants with altered sequence and function. In higher plants, AS has been implicated in a wide range of developmental and physiological processes including responses to stress [11-17]. Around 60% of Arabidopsis intron-containing genes undergo AS [20]. SFs are required for normal growth and development including control of flowering time, regulation of the circadian clock and stress responses suggesting that regulated AS of downstream targets is essential [10, 11, 21-24]. The importance of AS to the cold response has been demonstrated by altered cold sensitivity or tolerance when SFs are mis-expressed [10, 11, 24]. These studies strongly suggest that AS networks are central co-ordinators of the cold response. However, virtually nothing is known about the extent and timing of the contribution of AS or how transcription and AS determine the dynamic changes in the transcriptome required for response to cooling, acclimation, freezing tolerance, and survival.

RNA-sequencing provides the means to understand how AS impacts gene expression and transcriptome re-programming. However, the quality of AS analysis depends on accurate quantification of transcripts which in turn depends on mapping RNA-seq reads to the genome, construction of transcript isoforms and estimation of transcript levels from the number of aligned reads. Transcript assembly from short reads is often inaccurate and, for example, two of the best performing RNA-seq analysis programs, Cufflinks and StringTie, generate 35-50% false positives through mis-assembly of transcripts and non-assembly of *bona fide* transcripts both of which impact the accuracy of transcript quantification [25-27]. In addition, accuracy also decreases with increasing numbers of isoforms in a gene [26, 28]. Rapid, non-alignment programs, Salmon or kallisto, have high accuracy of transcript quantification [25, 26] but require well-annotated and comprehensive transcriptomes which are often limited in plant species. To improve the accuracy of transcript abundance and AS in RNA-seq analyses in Arabidopsis, we developed a comprehensive reference transcriptome dataset, AtRTD2 [27], for use with Salmon or kallisto. AtRTD2 contained over 82k transcripts giving greatly increased coverage of AS transcript isoforms than currently available in the Arabidopsis TAIR10 and Araport11 transcriptome datasets. Importantly, we demonstrated the increased accuracy of AS measurements using Salmon/AtRTD2 and the negative impact of missing transcripts [27]. Although AS has been detected in many Arabidopsis genes in response to stress conditions [11, 28, 29], the true scale and the dynamics of the AS response need to be addressed. To examine the dynamics of AS requires accurate quantification of individual transcripts in a time-series which takes full account of time-of-day variations in gene expression, due to photoperiod and circadian control.

Here, we exploit the enhanced ability of AtRTD2/Salmon to quantify transcript isoforms, the experimental design of the RNA-seq diel time series and robust analyses to capture the complexity and dynamics of changes in AS in response to lowering temperatures. The increased resolution from ultra-deep RNA-seq of multiple time-points identified 8,949 genes which were differentially expressed at the gene level (DE) and differentially alternatively spliced (DAS). These included 1,647 genes which were regulated only at the AS level, the majority of which had not been identified as cold-responsive by other methods. The high temporal resolution of both gene expression and AS shows the rapid and dynamic induction of both total gene expression and AS of thousands of genes in the first few hours after the onset of cold. Response to low temperature is thus controlled by genome-wide changes in both transcription and AS. Early cold response genes include specific splicing and transcription factors which undergo rapid AS that is sensitive to small changes in temperature. Our data suggest a mechanism whereby dynamic changes in AS of splicing regulators contribute to the changes in the transcriptome that condition the developmental and physiological processes required by plants to respond to constantly changing environmental conditions.

## Results

### Almost half of the expressed genes show cold-induced differential expression and/or differential alternative splicing

To examine changes in gene expression and AS in response to low temperature, we performed deep RNA-seq on a diel time-series of rosettes of 5-week-old Arabidopsis Col-0 plants grown at 20°C and transferred to 4°C (Fig. 1a). Rosettes were sampled at 3 h intervals for the last day at 20°C, the first day at 4°C and the fourth day at 4°C as described in Fig. 1a (see Methods). We generated over 360 million paired end reads for each of the 26 time-points and quantified transcript abundance in <u>t</u>ranscripts <u>p</u>er <u>m</u>illion (TPM) using Salmon [26] and AtRTD2-QUASI as reference transcriptome [27] allowing us to determine patterns of expression at both the gene level (sum of all transcripts of a gene) and at the individual transcript isoform level (Additional file 1: Figures S1 and S2). Principal component analysis of the gene level expression data from across the 26 time-points showed that temperature (32.4% of variance) and time of day (18.9% of variance) were the overwhelming drivers of gene expression (Fig. 1b). In order to analyse the time-series data at gene and transcript levels to obtain differential expression (DE), differential alternative splicing (DAS) and differential transcript usage (DTU) results, we developed an analysis pipeline that exploited the general linear models available in limma [30-32] (see Methods), which allowed biological repeats and time-of-day variation in expression to be taken into account in the statistical analysis to produce more accurate and robust results. It is important to note that in the time-series analysis, we compared gene and transcript abundances between equivalent time-points at 20°C and those in day 1 and day 4 at 4°C to remove the effects of time-of-day so that the changes detected are due to reduction in temperature (contrast groups in Additional file 1: Figures S1 and S2). We firstly analysed differential expression at the gene level (DE). A gene was considered differentially expressed if it showed a log_2_-fold change ≥1 (≥2-fold change) in expression in at least two consecutive contrast groups (adjusted p<0.01) (Fig. 2a; Additional file 1: Figure S1; Additional file 2: Table S1). Using these stringent criteria, we identified a total of 7,302 genes which were significantly differentially expressed in response to low temperature when compared to 20°C levels. Of these, 48.2% were up-regulated and 51.8% down-regulated (Additional file 1: Figure S3; Additional file 3: Table S2).

**Figure 1.**
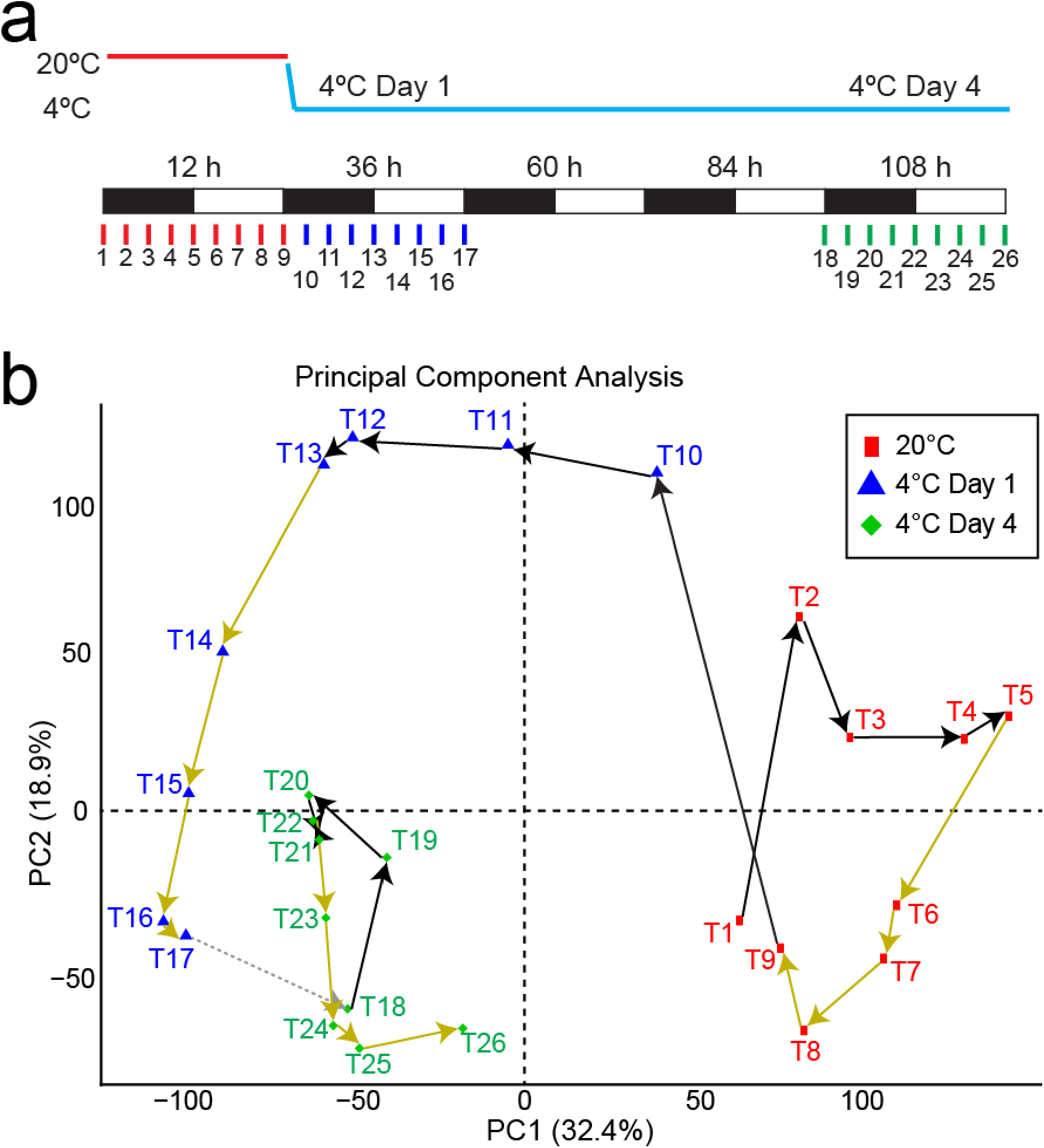
Analyses of the Arabidopsis response to low temperature in diel conditions. **a** Schematic representation of sampling strategy. Time-points of sampling are marked by vertical coloured lines and labelled from 1 to 26. Five-week-old Arabidopsis rosettes were harvested every 3 h over a 24 h-period at 20°C (red lines). At dusk, the temperature was gradually reduced to 4°C, and harvesting continued during the first day at 4°C (blue lines) and the fourth day at 4°C (green lines). Black boxes, 12 h dark; white boxes, 12 h light. **b** Principal Component Analysis of the Arabidopsis rosette transcriptome before and after a shift from 20°C to 4°C. Each data point (T1 to T26) refers to one time-point and represents the average gene expression (*n* = 3) from the RNA-seq data. The data points are connected by arrows in chronological order: black for 3 h of dark, yellow for 3 h of light. The dotted grey line joining T17 and T18 represents days 2 and 3 at 4°C. The first and second principal components (PC1 and PC2) account for more than 50% of the variance in the experiment. The cyclical arrangement of the data points – including segregation of light and dark-time samples – and their gradual shift upon cold treatment confirms that responses to photoperiod/circadian clock and temperature are the overwhelming drivers of gene expression differences.

**Figure 2.**
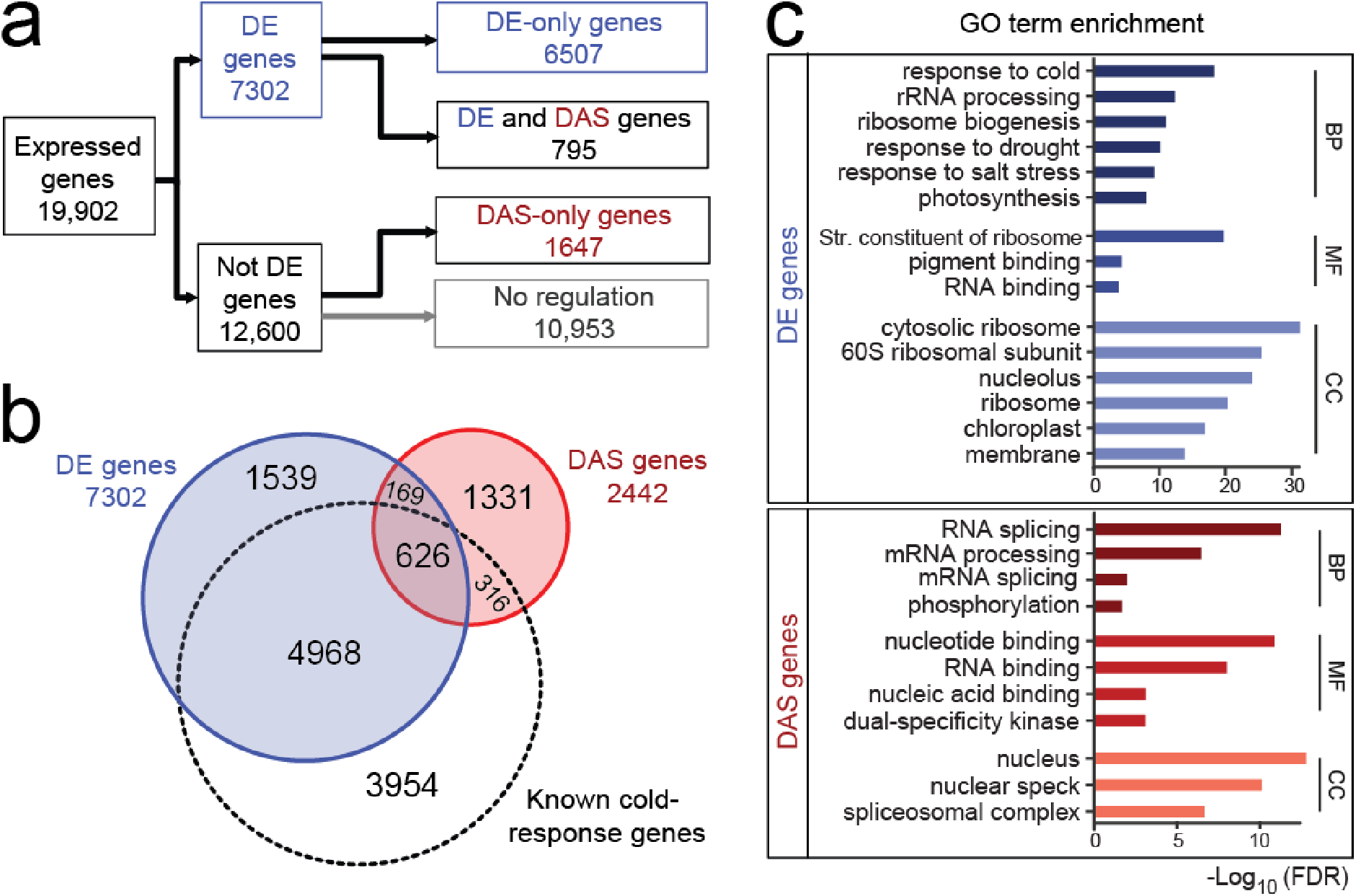
Differential Expression (DE) and Differential Alternative Splicing (DAS) analyses of Arabidopsis response to low temperature. **a** Flow chart showing the distribution of the 8,949 DE (blue) and DAS (red) genes (Additional file 2: Table S1). The DE and DAS gene sets are largely different with only 795 (11.26%) in common (overlap between blue and red circles in b). **b** Euler diagram of DE (left) and DAS (right) genes identified here and compared to known cold-response DE/DAS genes (dashed circle). Information on known cold-response DE/DAS genes can be found in Additional files 4 and 5: Table S3 and S4, respectively. **c** Most significantly enriched Gene Ontology (GO) terms for DE (shades of blue) and DAS (shades of red) genes. Bar plots of –log_10_ transformed FDR values are shown. BP: biological process; MF: molecular function; CC: cellular component.

Secondly, we used the transcript-level data to identify genes that were differentially alternatively spliced (DAS) between contrast groups. F-tests were carried out to examine the consistency of expression changes among the transcripts and gene (see Methods) to detect DAS genes. Criteria for genes being DAS were that in at least two consecutive contrast groups 1) at least one of the transcripts differed significantly from the gene with an adjusted p-value <0.01, and 2) at least one of the transcripts of the gene showed a **A** Percent Spliced (APS) of £0.1 (to keep genes where there is a significant change coming from transcript(s) with large differences in their relative abundance) (Fig. 2a; Additional file 1: Figure S2; Additional file 2: Table S1). We identified 2,442 DAS genes (Fig. 2a) of which 795 were also DE genes (regulated by both transcription and AS) and 1,647 genes that were not DE (regulated by AS only). Thus, of the total of 8,949 genes, which exhibited significantly altered levels of differential gene and/or transcript level expression, 27.3% were differentially alternatively spliced, consistent with widespread DAS in response to cold (Fig. 2a; Additional file 1: Figure S2; Additional file 2: Table S1). At any particular time-point, between ca. 600-3,700 genes were DE and ca. 400-1,450 genes were DAS when compared to 20°C levels (Additional file 1: Figure S3).

To pinpoint the individual transcripts responsible for a gene being identified as a DAS gene, a differential transcript usage (DTU) analysis was carried out by testing the expression changes of every transcript against the weighted average of all the other transcripts of the gene using a t-test (Additional file 1: Figure S2). DTU transcripts were identified as those which differed from the gene level and which showed a APS of >0.1 in at least two consecutive contrast groups with an adjusted p-value of <0.01. In total, 34% (4,105) of expressed transcripts of DAS genes were classed as DTU (Additional file 2: Table S1) of which ∼70% were protein-coding and ∼30% contained premature termination codons (PTC).

We next explored differences between immediate responses upon transfer to low temperature and the response to prolonged cold acclimation by comparing which genes were DE and DAS in day 1 and/or day 4 after transfer to 4°C. Around 50% (3,573 genes) and 60% (1,440) of DE and DAS genes, respectively, were common to both days with the remainder being either unique to day 1 or day 4 (Additional file 1: Figure S4; Additional file 2: Table S1). Thus, changes in gene-level expression and AS occurred throughout the cold period: either transiently (occurring in day 1 at 4°C and returning to 20°C levels by day 4), persisting throughout the cold period (occurring in day 1 at 4°C and remaining at day 4), or occurring later, only in day 4. We propose that these patterns of gene-level expression and AS reflect different contributions to low temperature perception, initial cold responses and physiological acclimation to cold and freezing temperatures.

### DE and DAS analyses identify novel cold response genes and AS of cold regulators

Previous analyses of differential gene expression in wild-type Arabidopsis plants exposed to cold used microarrays [12, 33, 34] and, more recently, RNA-seq [6, 7, 35] (Additional files 4 and 5: Table S3 and S4, respectively). There was a substantial overlap between the cold response DE genes identified in those studies and the DE genes identified here (Fig. 2b). Critically, we identified an additional 1,708 novel DE cold response genes (Additional file 6: Table S5). As expected, we showed cold induction of *CBFs* and selected *COR* genes (none of which undergo AS) (Additional file 1: Figure S5a and b). However, the first significant increase in *CBF* expression (*CBF2*) was detected between 3 and 6 h after onset of cold treatment (applied at dusk) whereas expression of *CBFs* has been detected within less than 1 h and peaked at around 1-2 h in other studies conducted in constant light [36]. The differences in timing of expression of the *CBF* and *COR* genes seen here will partially reflect variation in the age of plants tested and the experimental conditions (summarised in Additional file 7: Supplementary Notes) that here included the application of cold at dusk with gradual reduction in temperature at the beginning of the cold treatment (13.5°C at 30 min, 8°C at 1 h, 5.5°C at 90 min and 4°C at 2 h; Additional file 1: Figure S6). On the other hand, the differences in the detection of DE genes (Fig. 2b) most likely reflect the quality and depth of our data, including comparisons that remove the effects of the time-of-day variation, a much higher number of time-points, and robust statistical analysis (Additional file 4: Table S3; Additional file 7: Supplementary Notes).

Differential alternative splicing in response to cold was analysed previously using an algorithm to extract individual probe hybridisation data from microarrays of Arabidopsis seedlings exposed to 4°C [12]. Comparison with TAIR9 transcripts identified over 200 DAS transcripts although only half of those tested experimentally were validated [12]. While demonstrating that AS occurred in response to cold, the limited resolution of this approach only detected a fraction of DAS genes and transcripts. In comparison, the analysis here identified 2,442 DAS genes and 4,105 DTU transcripts. In particular, we identified 1,500 novel cold response DAS genes of which 1,331 displayed regulation only by AS (DAS-only) (Fig. 2b; Additional file 6: Table S5). Among the DAS genes identified here and which had previous evidence of involvement in the cold response, we observed dynamic changes in AS in, for example, *REGULATOR OF CBF EXPRESSION (RCF1)*, *PHYTOCHROME-INTERACTING FACTOR 7* (*PIF7*), *PHYTOCHROME B* (*PHYB*) and *SUPPRESSOR OF FRIGIDA 4* (*SUF4*) (Additional file 1: Figure S5c-f). *RCF1* encodes a cold-inducible RNA helicase required for cold tolerance [37]. It produces transcript isoforms which differ by AS of introns in the 3’ untranslated region (UTR) which may cause particular isoforms to be retained in the nucleus or trigger NMD to regulate RCF1 expression at different temperatures. PIF7 is a transcriptional repressor of the circadian expression of CBFs and is involved in photoperiodic control of the CBF cold acclimation pathway; its activity is regulated by the clock component, TOC1, and PHYB [38]. PIF7 shows temperature-dependent AS altering the relative levels of the fully spliced, protein-coding transcript and a non-protein-coding transcript with retention of intron 1. PHYB is a photoreceptor required for photomorphogenesis and is involved in the interaction of light- and cold-signalling pathways [39, 40]. The two *PHYB* transcript isoforms differ by removal or retention of intron 3 which alters the C-terminal sequences of the predicted proteins by the presence/absence of the histidine kinase related domain (HKRD) required for full functionality [41, 42]. Finally, SUF4 is required for delayed flowering in winter-annual Arabidopsis and is involved in chromatin modification of *FLOWERING LOCUS C* (*FLC*) [43]. *SUF4* AS transcripts differ by splicing/retention of intron 6 to alter the C-terminal sequences [44] and here, we demonstrate rapid cold-induced changes in AS (Additional file 1: Figure S5). Thus, in addition to the previously observed cold-induced changes in gene level expression, we demonstrate that low temperature-dependent AS is an important further layer of regulation that controls the abundance of cold response genes and transcripts.

### DE and DAS genes have different predicted functions

The DE and DAS gene sets were largely different with an overlap of only 795 genes (Fig. 2a). The most significantly enriched Gene Ontology (GO) terms for DE genes correlated with known physiological and molecular events in the cold response such as response to various stresses and increased ribosome production (Fig. 2c; Additional file 8: Table S6). Hierarchical clustering of total gene expression levels of DE genes revealed transient, adaptive and late expression profiles in response to cold and regulation between light and dark (Fig. 3; Additional file 1: Figures S7 and S8; Additional file 9: Table S7). Transcript expression profiles of individual DE genes showed similar responses (Additional file 1: Figure S9). Functional annotation of individual DE gene clusters was associated with response to cold, abscisic acid and drought (cluster 6), decreased photosynthesis (cluster 10), increased ribosome production (cluster 3) and membrane and lipid biosynthesis (cluster 12) (Additional file 1: Figures S7 and S8; Additional file 9: Table S7). Cluster 12 shows highly increased gene level expression in the first 3 h of cold treatment reflecting the reconfiguration of membranes in response cold to maintain fluidity and protect against subsequent freeze damage [3].

**Figure 3.**
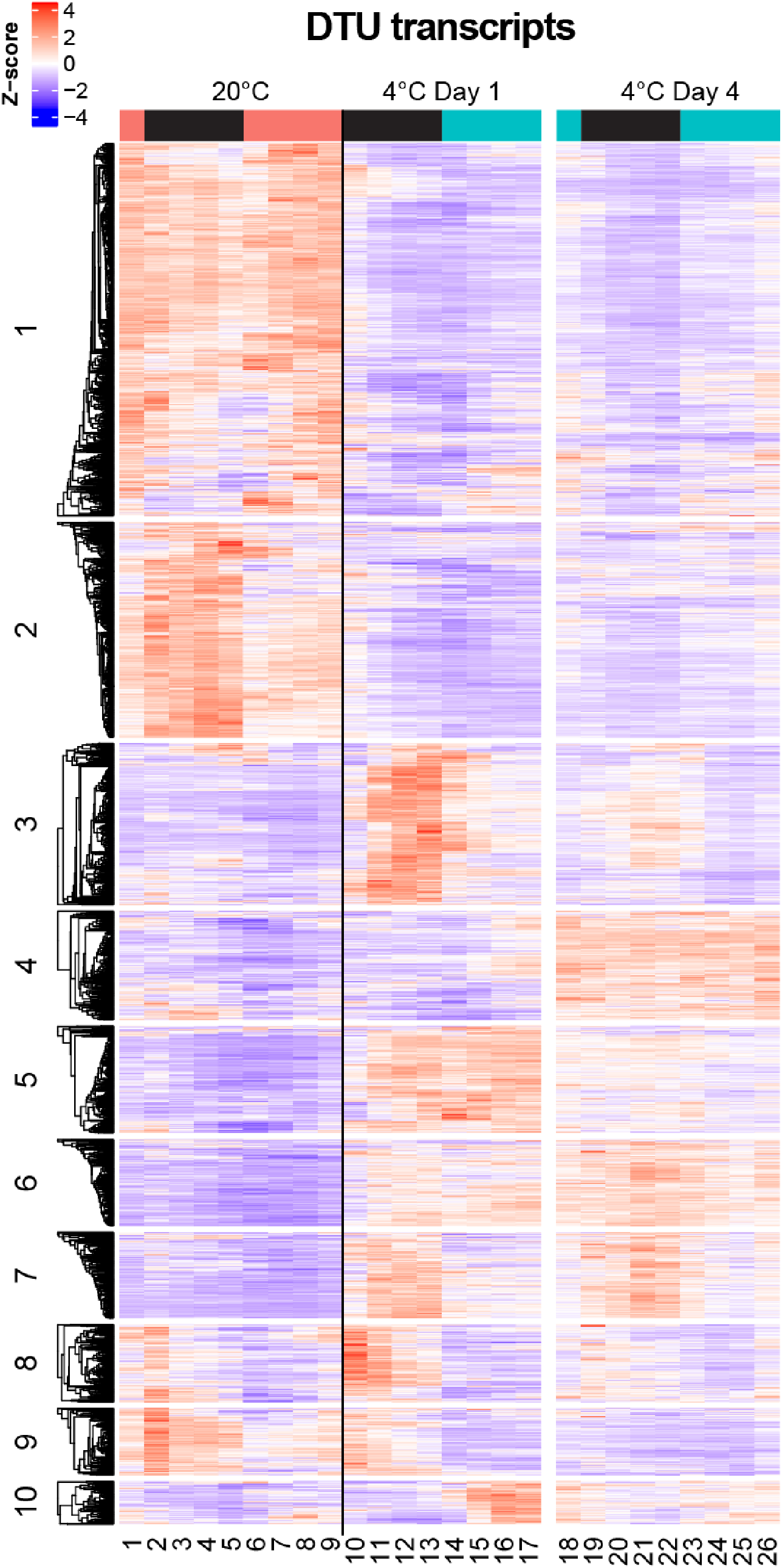
Heat map of DTU transcripts from DAS genes. DTU transcripts from DAS genes show segregation into 10 co-expressed clusters. For simplicity, transcripts that do not fall into any cluster have been removed from heat map (*n* = 36). Clusters 1 and 2 show transcripts down-regulated upon cold. Clusters 3, 5, and 10 show clear transient changes in AS isoform transcripts at different times during Day 1 at 4°C while cluster 4 *(n =* 326) shows late up-regulation of transcripts on the fourth day at 4°C. Clear gain in rhythmic expression of AS transcripts upon cold is seen in cluster 7 *(n =* 258). Cluster 8 *(n =* 233) includes transcripts with increased expression within the first 3 h of cold treatment. The *z*-score scale represents mean-subtracted regularized log-transformed TPMs. The coloured bars above the heat map indicate whether samples were exposed to light (coloured) or dark (black) in the 3 h before sampling.

For the DAS genes, the most enriched functional terms were related to mRNA splicing (Fig. 2c; Additional file 8: Table S6). One hundred and sixty-six (7%) DAS genes were RNA-binding proteins, spliceosomal proteins or SFs. Hierarchical clustering of the DTU transcripts and expression profiles of individual DAS genes also showed transient, adaptive and late expression patterns (Fig. 3; Additional file 1: Figures S10-12). Functional annotation of the genes in the individual DTU clusters showed enrichment of terms involved the plasma membrane and signal transduction (cluster 8, *p*<0.001) as well as regulation of transcription (cluster 3, *p*<0.001), RNA splicing (cluster 5, *p*<0.0001) and chromatin binding (cluster 1, *p*<0.0001) (Fig. 3).

### Rapid cold-induced changes in AS accompany the major transcriptional responses

The high temporal resolution of the time-course allowed us to examine the relative dynamics of the DE and DAS changes. The statistical model used in this analysis allowed us to determine precisely at which time-point each DE and DAS gene first showed a significant change (start time-point), along with the magnitude and duration of that change. The dynamics of the changes of DE and DAS genes was compared by plotting the distribution of start time-points (Fig. 4a, b). DE and DAS genes peaked at 6-9 h after onset of cold (Fig. 1a, T11 and T12) and 6 h after onset of cold (Fig. 1a, T11), respectively. 62.2% (1,520) of the DAS genes and 47.6% (3,473) of the DE genes were detected within the first 9 h of low temperature (Fig. 4a, b; Additional file 10: Table S8). The speed of induction is highlighted by 648 and 388 genes showing significant DE and DAS after only 3 h of cold (T10) (Fig. 4a, b), respectively, despite the gradual reduction in temperature which takes 2 h to reach 4°C (Additional file 1: Figure S6). Notably, three-quarters (76.5%; 849) of the DAS genes detected within 9 h of cold (Fig. 1a, T10-T12) also had large AS changes with at least one transcript having ΔPS >0.2 (Additional file 11: Table S9). A further indicator of the speed of AS response and the sensitivity of some AS to reductions in temperature was demonstrated by identifying those DAS genes which show isoform switches (IS), where the relative abundance of different isoforms is reversed in response to cold (Additional file 1: Figures S11 and S12). We developed the Time-Series Isoform Switch (TSIS) program to identify ISs in time-series data [45]. Using TSIS, a total of 892 significant (*p*<0.001) ISs that involved two abundant transcript isoforms (see Methods) were identified in 475 unique DAS genes (Fig. 5a; Additional file 12: Table S10). The ISs involved different types of AS events and 77% involved at least one protein-coding transcript isoform (Fig. 5a, b; Additional file 12: Table S10). These either generated isoforms which coded for different protein variants, or where AS occurred in the 5’ or 3’UTR, the transcript isoforms coded for the same protein, or one of the transcripts was non-protein-coding (e.g. PTC-containing). TSIS determines the two time-points between which a significant isoform switch occurs and, consistent with the rapid changes in AS, the majority (57%) occurred between 0-6 h following transfer to 4°C (Fig. 5a; Additional file 12: Table S10). Thus, immediately in response to lowering temperature, there are waves of transcriptional and AS activity involving thousands of genes.

**Figure 4.**
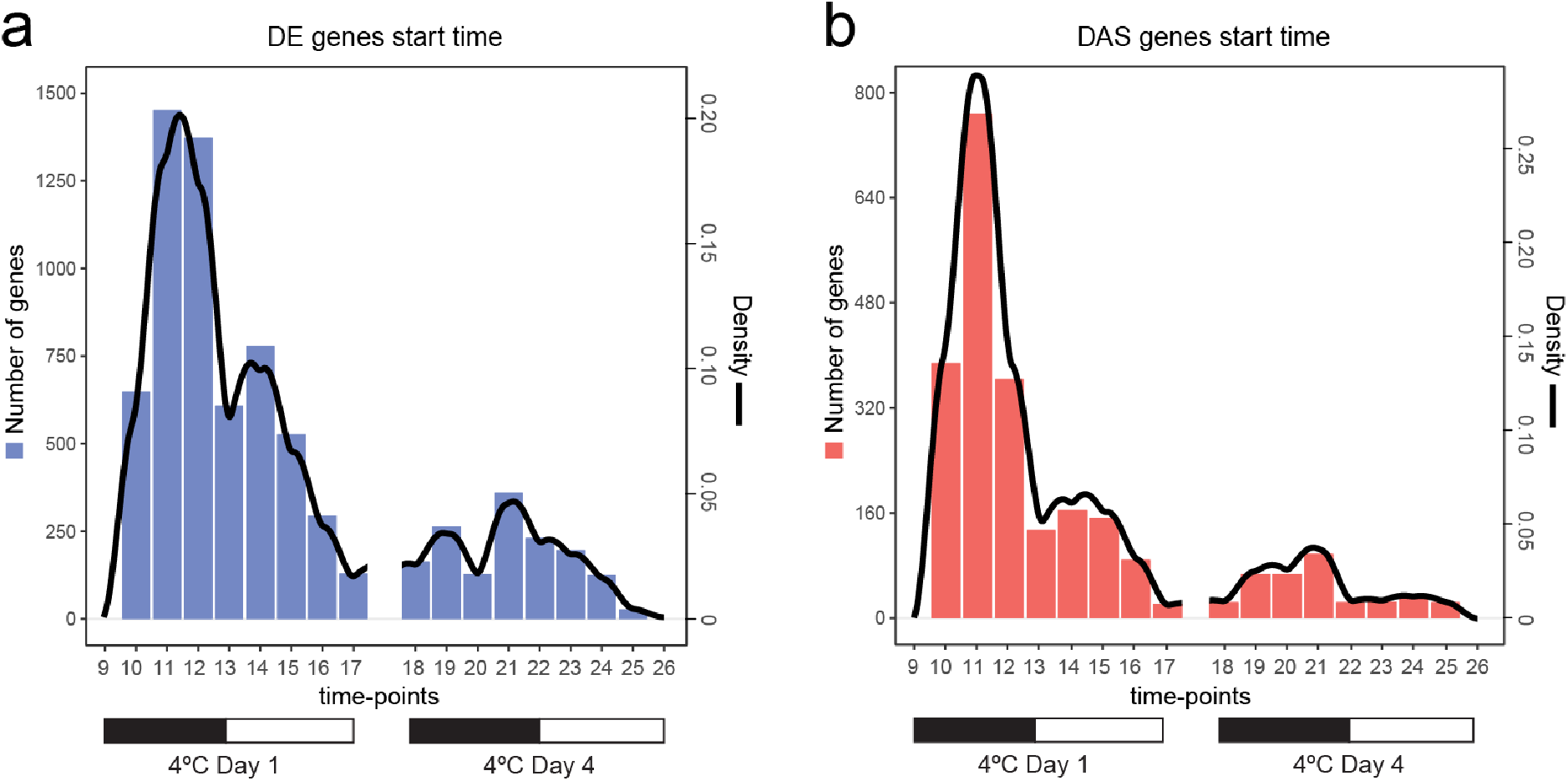
Rapid changes in DE and DAS genes in response to cold. Histograms and density plots of the time-points at which the **a** 7,302 DE and **b** 2,442 DAS genes first become significantly different in Day 1 and Day 4 at 4°C compared to 20°C. The genes that first show significant differences after longer exposure to cold (Day 4 at 4°C) represent 20.45% of DE and 14.73% of DAS genes. Each gene is represented only once in each histogram (left y-axis). The estimated density line of the number of genes illustrates the early waves of transcriptional and alternative splicing responses (right y-axis).

**Figure 5.**
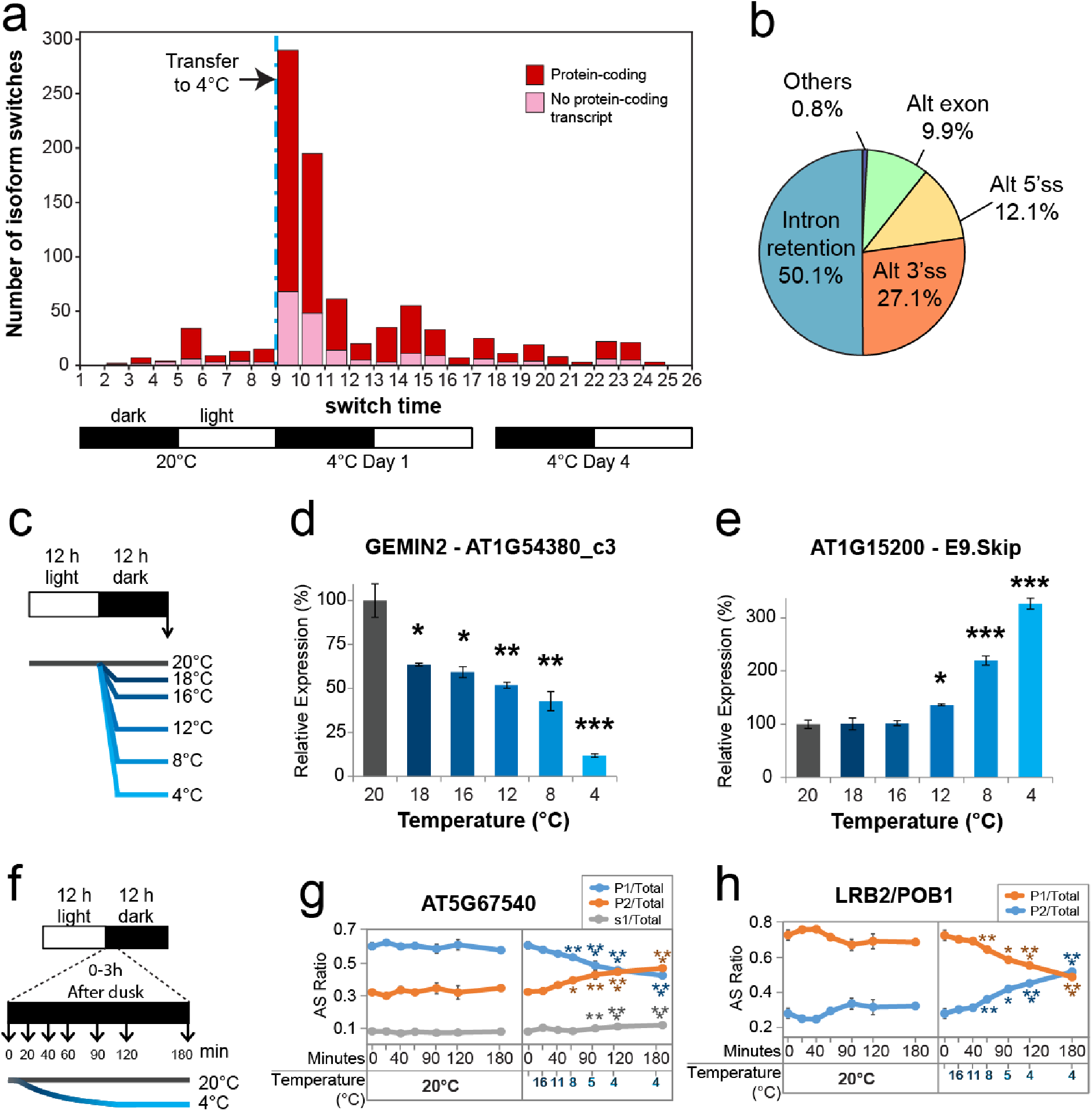
Sensitivity of AS to low temperatures. **a** Frequency over time of isoform switches in the RNA-seq time-course. Each isoform switch involved “abundant” transcripts, (i.e. expression of each transcript makes up at least 20% of the total expression of the gene in at least one time-point). Proportion of protein-coding transcripts is also shown and represents either production of different protein-coding isoforms or transcripts encoding the same protein where key AS events are in the UTR region. Data between T17-T18 represent ISs that occurred between Day 1 and Day 4. **b** Proportion of the major types of AS events involved in isoform switches in **a** was measured with SUPPA [46]. **c** Experimental design for assessing long-term changes in AS induced by small reductions in temperature initiated at dusk. Sampling of 5-week-old Arabidopsis rosettes occurred at dawn, after 12 h of temperature reduction, and is marked by a vertical arrow. **d** AS of novel cold-response gene GEMIN2 (PTC-containing transcript AT1G54380_c3) is sensitive to reductions in temperature of 2°C. **e** Exon 9 skipping (E9.Skip) of novel cold-response gene AT1G15200 (transcripts AT1G15200.1, AT1G15200_JS1, AT1G15200_JS2) is sensitive to reductions in temperature of 8°C. **f** Experimental design for assessing immediate changes in AS induced by gradual reductions in temperature initiated at dusk. Sampling of 5-week-old Arabidopsis rosettes occurred at different time-points after dusk and is marked by vertical arrows. **g** AS of novel cold-response gene AT5G67540 (transcripts AT5G67540_P1, AT5G67540_P2 and AT5G67540_s1) is affected within 1 h of gradual reduction in temperature. **h** AS of novel cold-response gene *LIGHT-RESPONSE BTB 2* (LRB2/POB1, transcripts AT3G61600_P1, AT3G61600_P2) is affected within 1 h of gradual reduction in temperature. In **d-e** and **g-h**, Tukey t-tests were performed to compare each temperature reduction results against 20°C control. Significant differences are labelled with asterisks (*, *p*<0.05; **, *p*<0.01; ***, *p*<0.001).

### Cold-regulated expression and AS of transcription and splicing factors

The waves of differential expression and AS in response to cold likely reflect regulation by transcription factors (TFs) and splicing factors/RNA-binding proteins (SF-RBPs). Differential expression of TFs in response to cold has been well documented (see Background). Here, 532 of the 2,534 TFs predicted in Arabidopsis (Additional file 13: Table S11) were significantly regulated only at the gene level (DE-only) (Table 1). However, a third (271) of TFs with cold-induced expression changes involved AS (Table 1), which potentially affects the levels or function of TF proteins. Similarly, of the 798 predicted SF-RBP genes (see Methods; Additional file 13: Table S11), 197 were DE-only, 33 DE+DAS and 133 DAS-only (Table 1). Thus, many TF and SF-RBP genes were regulated by AS in response to lower temperatures. The majority have not previously been associated with the cold response and represent putative novel cold response factors (Additional file 14: Table S12). We next identified the TF and SF-RBP genes with the fastest (0-6 h after onset of cold) and largest changes in expression and AS (log_2_ fold change ≥1.5 (equivalent to 2.83-fold change) for DE genes and ΔPS >0.25 for at least one transcript in DAS genes) (Additional file 14: Table S12). Fifty-nine TF and 47 SF-RBP DAS genes were identified as “early AS” genes. The TF genes included a high proportion of circadian clock genes as well as genes associated with abiotic stress, flowering time and hormone responses (Additional file 15: Table S13). The SF-RBP genes included serine-arginine-rich (SR) and heterogeneous ribonucleoprotein particle (hnRNP) protein genes known to be involved in stress responses and regulation of the clock. For many of the early AS genes, the AS changes were either only observed in day 1 at 4°C, or persisted through the cold treatment, and many involved isoform switches (Additional file 15: Table S13). Thus, both transcription and AS determine expression levels of TFs and SFs and many of these genes were regulated rapidly in response to reduction in temperature.

**Table 1.** Splicing factor/RNA-binding protein and transcription factor genes that are differentially expressed (DE) and/or differentially alternatively spliced (DAS) in response to lowering of temperature.

|  | DE-only | DE+DAS | DAS-only | Total |
| --- | --- | --- | --- | --- |
| Total genes | 6507 | 795 | 1647 | 8949 |
| SF-RBP (798) | 197 | 33 | 133 | 363 |
| TFs (2534) | 532 | 85 | 186 | 803 |

### Speed and sensitivity of AS responses to small reductions in temperature

The rapid and large changes in AS suggest that many AS events are sensitive to relatively small changes in temperature. To investigate further, we examined the effect on AS of early AS genes when the temperature was lowered from 20°C to 18°C, 16°C, 12°C, 8°C and 4°C for 12 h (Fig. 5c-e; Additional file 1: Figure S13). We observed that the relative levels of AS transcript isoforms were dependent on the temperature. Three of the six genes analysed showed significant changes in the level of at least one AS isoform with only a 2°C reduction in temperature (to 18°C) while others were affected by 4°C or 8°C reductions (Fig. 5e; Additional file 1: Figure S13). We then examined the speed and sensitivity of AS by taking multiple samples between 0 and 3 h after the onset of cold (Fig. 5f-h; Additional file 1: Figure S14). Of eight genes examined, one showed significant AS changes within 40 minutes when the temperature had reached 11°C, five within 60 min (at 8°C) and two within 180 minutes (at 4°C) (Fig. 5g-h; Additional file 1: Figure S14).

### *U2B”-LIKE* is regulated at the AS level and is required for freezing tolerance

Many of the early AS genes, including TFs and SFs, showed large and rapid changes in AS which alter the levels of protein-coding transcripts (Additional file 15: Table S13). We hypothesised that such significant changes in AS in response to low temperature are important in the overall cold acclimation process of the plant and lead to improved ability to tolerate freezing conditions after acclimation. In support of this, four of the early AS genes have previously been shown to be required for cold acclimation and tolerance to freezing: *RCF1* and *STA1* [37], *GEMIN2* [22] and the LAMMER kinase, *AME3* [47] (Table 2). To examine whether other early AS genes may be involved in cold acclimation, we selected the SF-RBP gene *U2B”-LIKE* because it was a novel DAS-only gene with an adaptive expression pattern. We isolated a knockout mutant of the *U2B”-LIKE* gene (AT1G06960; Fig. 6a; Additional file 1: Figure S15). *U2B”-LIKE* has two main AS transcripts, the fully spliced protein-coding mRNA and an isoform with retention of intron 4 (I4R) (P1 and P2, respectively - Fig. 6a). In wild-type plants, the protein-coding P1 transcript isoform showed rhythmic expression at 20°C, loss of rhythm during day 1 at 4°C, maintaining a high level of expression throughout the remaining cold treatment (Fig. 6a). In freezing tolerance tests conducted at −8.0°C and −8.5°C, the *u2b”-like* mutant plants showed greater sensitivity to freezing; *u2b”-like* did not survive freezing at −8.5°C after cold acclimation while wild-type plants recovered (Fig. 6b). *u2b”-like* mutant and WT plants both recovered at −8.0°C. Differential sensitivity of the mutant was confirmed in quantitative electrolyte leakage analyses; leaf tissue of *u2b”-like* suffered significantly increased ion leakage (cellular damage) at −10°C than wild type plants (Fig. 6c) indicating that expression of *U2B”-LIKE* is required for cold acclimation and freezing tolerance.

**Figure 6.**
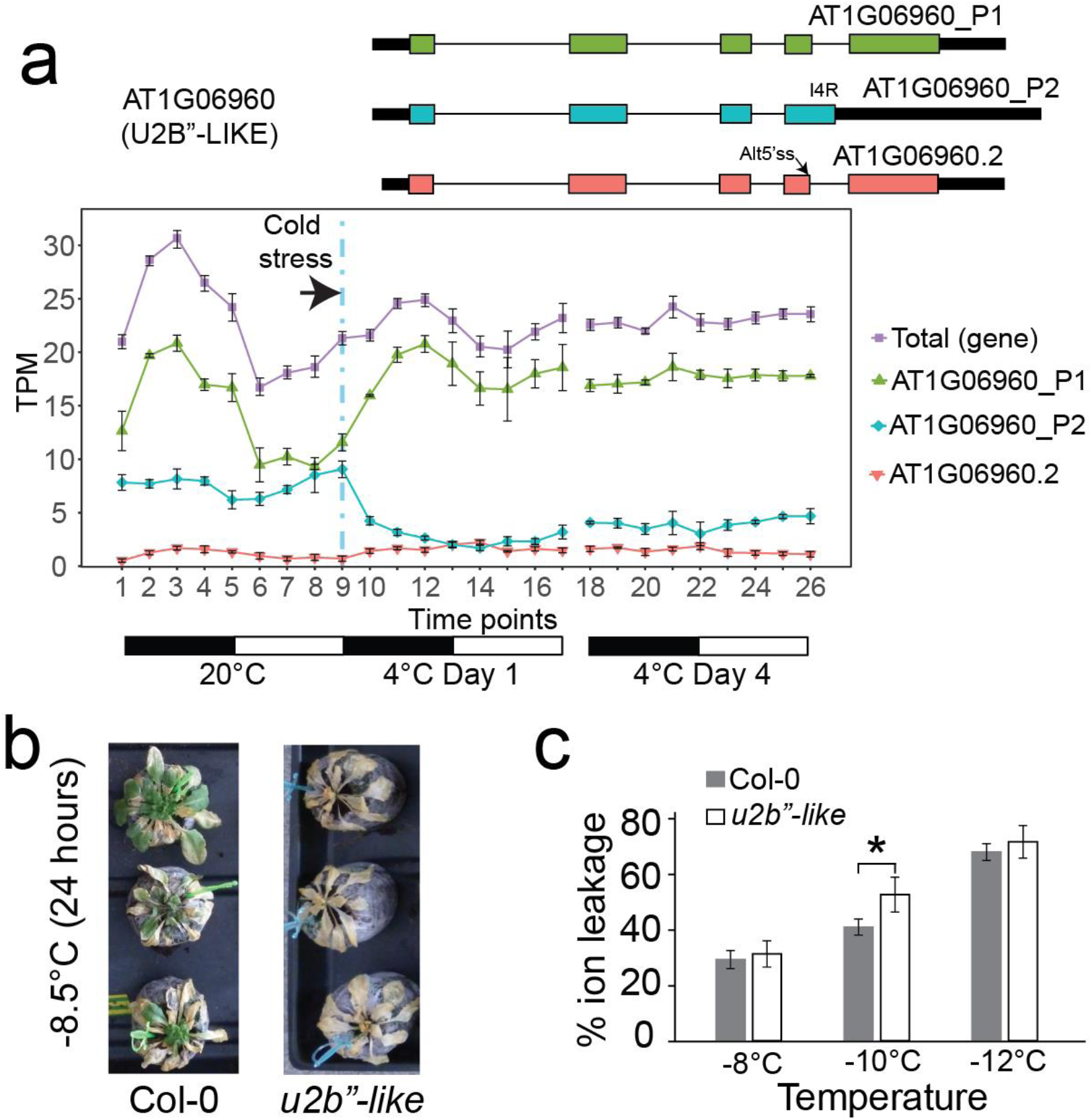
AT1G06960 (*U2B”-LIKE*) expression profile and freezing assays of knock-out line. **a** Structures of highly expressed *U2B”-LIKE* transcripts (black boxes, UTR; colour-coded boxes, exon CDS) and gene/transcript expression profile across the time-course. I4R: Intron 4 Retention; Alt5’ss, alternative 5’ splice site. Black:white bars below expression plots represent 12 h dark:light cycles. **b** Freezing sensitivity of cold-acclimated Col-0 and *u2b”-like* mutants showing recovery of wild-type and non-recovery of *u2b”-like* mutant plants at −8.5°C. **c** Cellular ion leakage in Col-0 (WT) and *u2b”-like* (knock-out mutant) leaf discs subjected to different freezing temperatures before thawing (*n* = 4). Transformed ion leakage data were used in a one-tailed t-test which confirmed *u2b”-like* loses more electrolyte than wild-type Col-0 at −10°C (*p* = 0.0263, represented by *). Each bar of the plot represents average ion leakage values. In **a** and **c**, error bars are standard error of the mean. In **b** and **c** plants were grown at 20°C for 4-5 weeks and cold acclimated at 5°C for 2 weeks before freezing assay.

**Table 2.** DAS-only SF-RBP genes with established role in freezing tolerance/acclimation.

| Gene name | Gene ID | Maximum ΔPS value at 0-6 h after cold | Reference |
| --- | --- | --- | --- |
| RCF1 | AT1G20920 | ΔPS >0.3 | [37] |
| STA1 | AT4G03430 | ΔPS >0.24 | [37] |
| GEMIN2 | AT1G54380 | ΔPS >0.3 | [22] |
| AME3 | AT4G32660 | ΔPS >0.25 | [47] |

Arabidopsis contains two *U2B”*-related genes: *U2B”* (AT2G30260) and *U2B’’-LIKE* (AT1G06960). The two proteins are very similar: 80% identical and 90% similar at the protein level (Additional file 1: Figure S16). U2B” is an U2snRNP-specific protein which binds, along with U2A’, to stem-loop IV of U2snRNA in both plants and human (Additional file 7: Supplementary Notes). In the *u2b”-like* mutant, there was no expression of *U2B”-LIKE* but expression of the *U2B”* paralogue (which was neither DE nor DAS in cold) (Additional file 1: Figure S15) was detected, suggesting that *U2B”* protein could not compensate for the lack of U2B”-LIKE in the *u2B”-like* mutant and therefore that they had functionally diverged. To investigate whether *U2B”-LIKE* affected AS regulation, we then compared AS patterns of 41 genes (including 34 DAS or DE+DAS genes identified here) in wild-type and *u2b”-like* mutant plants. Five genes showed significantly different AS (*p*<0.05 and >10% difference in splicing ratio between the mutant and wild-type) (Additional file 1: Figure S17; Additional file 16: Table S14). These included decreased levels of fully spliced, protein-coding transcripts of *PIF7* (AT5G61270; Additional file 1: Figure S17) which along with TOC1 and PHYB represses expression of CBFs [40] and *HOMOLOGUE OF HY5* (*HYH*), a clock input gene. Thus, U2B”-LIKE is one splicing factor that contributes to correct splicing of *PIF7* linking U2B”-LIKE-dependent AS to regulation of the major cold response pathway. Therefore, the freezing sensitivity of the *u2b”-like* mutant may be due to altered AS and expression of specific genes required for cold acclimation (Additional file 7: Supplementary Notes).

## Discussion

Dynamic changes in expression at both the gene and transcript/AS levels occur in Arabidopsis plants in the process of cold acclimation. The use of the new AtRTD2 transcriptome of Arabidopsis with Salmon, the high-resolution time-course with a statistical model that takes into account time-of-day variations, and the novel analysis methods have captured a much higher degree of complexity of regulation in response to cold. In particular, we demonstrate the dynamic contribution of AS by the rapid cold-induced wave of AS activity accompanying the transcriptional response (Fig. 4) and the sensitivity of AS of some genes to small reductions in temperature (Fig. 5). We also significantly demonstrate the extent of AS by showing that over 2,400 genes are regulated by AS in response to cold with over 1,600 regulated only at the AS level (Fig. 2). The massive changes in expression and AS involved thousands of genes reflecting activation of both transcription and splicing pathways and networks. The speed and extent of the cold-induced AS suggest that AS, along with the transcriptional response, is a major driver of transcriptome reprogramming for cold acclimation and freezing tolerance.

With over 2,400 genes regulated by AS, multiple different mechanisms are likely to control the splicing decisions. Reduction in temperature to 4°C is expected to reduce the rate of biochemical reactions and potentially affect transcription and splicing. We observed that the vast majority of introns in the pre-mRNAs of all the cold-expressed genes are efficiently spliced throughout the cold treatment. Therefore, low temperature does not cause a general defect in splicing reflecting the ability of temperate plants to grow in a wide range of fluctuating temperatures. Nevertheless, low temperatures may directly affect AS regulation. For example, in mammals, secondary structures in pre-mRNAs affect splice site selection [48] and cooling could stabilise such structures. Similarly, splicing is largely co-transcriptional and slower rates of RNA polymerase II (PolII) elongation promote selection of alternative splice sites [49]. Both of these mechanisms will undoubtedly be involved in the cold-induced AS changes of some of the genes seen here. However, the sensitivity of AS to reductions in temperature of only a few degrees and clear rhythmic expression profiles of AS transcript isoforms in plants exposed to constant 4°C temperature for four days (e.g. cluster 7 in Fig. 3 and Additional file 1: Figure S10) argue against such mechanisms being widely responsible for the cold-induced AS changes observed here. Local or global DNA methylation and chromatin modifications can also affect the rate of PolII elongation or help to recruit SFs to affect splice site selection [49]. In plants, epigenetic regulation is responsible for suppression of *FLC* by vernalisation in the seasonal response to cold [50]. Furthermore, altered histone 3 lysine 36 tri-methylation (H3K36me3) was recently shown to affect some AS events induced by higher ambient temperatures within 24 h [14]. Alongside dynamic changes in histone marks at specific stress-induced genes [51, 52] it is likely that some of the cold-induced AS here reflects local epigenetic changes.

We showed that the levels of hundreds of TF and SF-RBP gene transcripts changed in response to cold at both the transcriptional and AS levels. Therefore, splicing decisions in the physiological response to low temperature are most likely controlled by altered abundance, activity or localisation of SFs or other RNA-interacting proteins [11, 18, 19, 24, 53]. In particular, we identified TF and SF-RBP genes with large and rapid changes in AS. Most of the early AS transcription factor genes were regulated only by AS and, therefore, had not been identified previously as cold response transcription factors. Nevertheless, the rapid cold-induced changes in the AS of some known cold response genes: *CAMTA3* which activates the CBFs [54], and *VRN2* and *SUF4* which are involved in vernalisation and silencing of *FLC* [43, 50], have not been described previously and our results introduce AS as a novel component in their regulation. It will be interesting to address the function of the novel AS-regulated TFs and the function of AS of these and known cold response TFs on cold acclimation and vernalisation in future experiments.

In contrast, the early AS SF-RBP genes included SR and hnRNP protein genes known to respond to changes in temperature (e.g. *SR30*, *RS40*, *GRP8*, *SR45A*, *PTB1*, *RBP25* etc.) [11, 23, 24]. Many SF-RBP genes are regulated by AS-NMD and the rapid induction of AS in these and other early AS genes affects the abundance of protein-coding transcripts and presumably of the splicing factors themselves to alter AS of downstream targets. Various spliceosomal and snRNP protein genes are also among the early AS genes. These include *GEMIN2* (snRNP assembly) which is cold-induced, involved in regulation of the circadian clock, and enhances U1snRNP assembly to compensate for reduced functionality of U1snRNP at low temperatures [22]. Interestingly, a number of U1snRNP core and associated protein genes (*U1-70k*, *LUC7B*, *LUC7RL*, *PRP39A*, *RBM25*) [55, 56] respond rapidly to cold via AS. The early AS genes also include two wound-induced RNA-binding proteins, UBA2a and UBA2c [57] and may also therefore be involved in the cold response. Three LAMMER kinase genes (*AFC1, AFC2 and AME3*) which regulate SR proteins via phosphorylation showed changes in their expression due to AS suggesting that lower temperatures affect activation/deactivation of specific splicing factors which are targets of these kinases [47, 58]. In addition, over 20 putative RNA-binding proteins, kinases and RNA helicases with little or no known function are among the novel early AS genes. Four of the early AS SF-RBP genes (*RCF1*, *STA1*, *GEMIN2* and *AME3*; Table 2) have been shown to be involved in freezing tolerance [22, 47] and we provide initial evidence for another early AS gene, *U2B”-LIKE*, being involved in freezing tolerance and acclimation (Fig. 6; Additional file 7: Supplementary Notes). Our results identify over 100 splicing and transcription regulatory genes, whose expression is rapidly and drastically altered by AS in response to cooling. Future work will address the function of these putative regulators and specific transcript isoforms in cold acclimation.

The speed of change of AS may be one of the earliest responses to cooling. We showed significant AS within only 40-60 minutes of cooling and with subtle reductions in temperature of as little as 2°C (Fig. 5). Similar responses are seen in mammals where neuronal stimulation and rapid changes of intracellular sterols also activate splicing/AS within minutes without *de novo* transcription or protein production [59, 60]. In addition, a 1°C change in body temperature activated a program of AS changes within 30 min which involved temperature-sensitive phosphorylation of SR proteins and AS of the U2-associated factor, U2AF26 [61]. Therefore, cold-induced AS programs may similarly involve rapid phosphorylation/dephosphorylation of SFs such as SR proteins to modulate AS [61, 62]. Interestingly, the expression of a third of the early AS SF-RBP genes revealed here are also affected by increased ambient temperatures [63] and these genes may represent key temperature-dependent splicing regulators. Master splicing regulators have been postulated to drive splicing networks during cellular differentiation in mammals where regulatory modules of SF-RBPs and/or TFs establish and maintain new expression patterns [64, 65]. Auto- and cross-regulatory modules of some plant SFs are well-documented and may be important components of splicing networks [11, 24].

The cold response pathway in Arabidopsis involves Ca^2+^-dependent and MAP kinase signalling cascades which affect ICE1 phosphorylation, lead to activation of CBFs and other TFs and expression of COR genes [3, 66-68]. In animals, signalling pathways including calcium-dependent signalling control of phosphorylation and AS of specific SFs to regulate AS of downstream targets [62, 69]. In plants, stress signals affect both the phosphorylation status and sub-cellular localization of some SR proteins [70, 71] and sixteen of the early AS SF-RBPs have been shown to be phosphorylated in Arabidopsis cells [71]. The cold-induced waves of transcription and AS, the rapid AS responses of SF-RBPs and their potential for phosphorylation suggest a model where cold signalling pathways modulate both transcription and splicing factor levels and activity (Fig. 7). These regulatory factors, in turn, drive gene and splicing networks required to determine the overall reprogramming of the transcriptome for cold acclimation and freezing tolerance. These networks are reflected in dynamic changes in the DE and DAS gene sets that we observe across the time-series.

**Figure 7.**
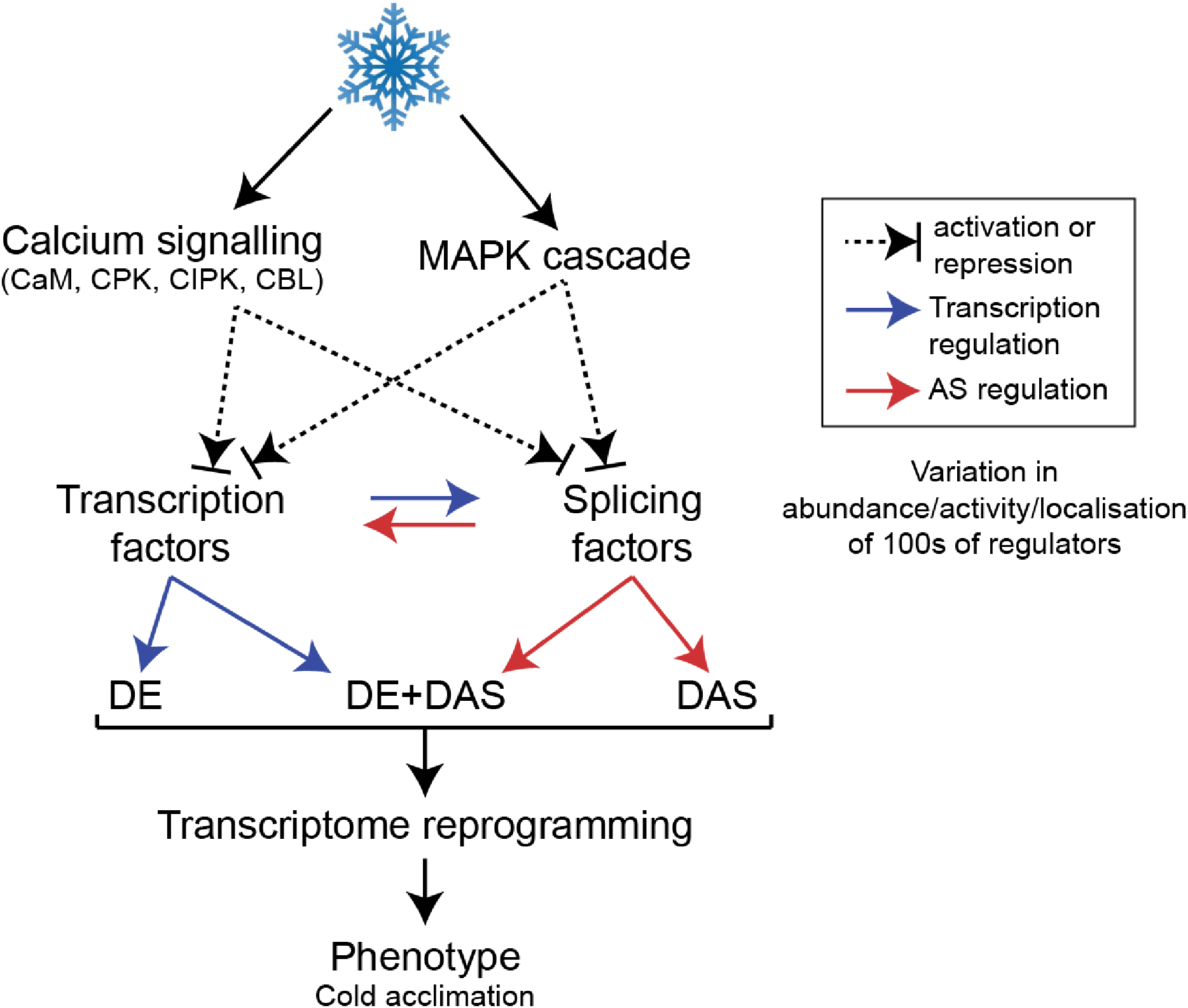
Model for the cold signalling pathway and its regulation of genome-wide gene expression. Cooling activates Ca^2+^-dependent kinases and MAP kinases which activate or repress transcription factors or splicing factors. These in turn regulate the transcription or AS of downstream genes including other TFs and SFs, thereby driving cascades of cold-induced gene expression.

Plants are exposed to a variety of temperature conditions. They require flexible regulatory systems that modify expression quickly and reversibly upon perception of constantly fluctuating temperatures throughout the day and night and during the regular 24 h cycle of warmer daytime and cooler night-time temperatures. They must also re-programme the transcriptome in cold conditions to allow the plant to acclimate, avoid freezing and survive as the intensity and duration of reduced temperatures increase (seasonal changes). The dynamic AS response, sensitivity of AS to small changes in temperature and the different behaviour of AS genes, where changes are transient or persist, demonstrated here, suggest that AS provides a level of flexibility to contribute to different stages of the progression from perception to acclimation. In particular, the speed of AS reactions may contribute to temperature perception by altering the activity of key TFs and SF-RBPs while the transcriptional response is being activated as seen in animal cells [60]. Such control could fine-tune expression of specific genes and pathways throughout the day as temperatures fluctuate. The rapid waves of transcriptional and AS activity within the first few hours of cold exposure, which include transcriptional activation of CBFs, other transcriptional response pathways and altered expression/AS of clock components may be involved in initial cold responses and in the normal 24 h day/night temperature cycle. Significant changes in expression/AS of many genes occur rapidly in the first day in the cold and persist throughout the cold period and other genes are only activated or repressed by transcription or AS after prolonged cold treatment (day 4); these genes may be important for establishing and stabilising changes in the transcriptome for acclimation. Thus, temperature-dependent AS is a mechanism to transduce temperature change signals into changes in expression. Dynamic and extensive changes in AS are also likely to drive plant responses to other abiotic stresses, to pests and diseases and in developmental programs alongside transcriptional responses. Construction of splicing and transcriptional networks from the data here will further define the contribution of AS, as an additional layer of regulation, and the interplay and co-ordination of the transcriptional and AS responses.

## Conclusions

The temporal resolution of the time-course of Arabidopsis plants exposed to low temperatures combined with enhanced diversity of AS transcripts in AtRTD2 and robust RNA-seq analyses has demonstrated the scale and dynamics of AS. Our results show the dynamic response of AS involving thousands of genes such that AS, alongside transcription, is a major component of the transcriptomic response to cold. By identifying hundreds of novel AS-regulated, cold-responsive genes including transcription and splicing factor genes, the complexity of regulation of expression has been significantly increased. Taken with rapid and sensitive changes in AS, transcriptional and AS activity may provide the flexibility to allow plants to adapt to changes in temperature over different time-scales and survive extremes. It is likely that other abiotic and biotic stresses will also include major changes in AS as part of transcriptome reprogramming.

## Methods

### Plant Material and Growth Conditions

*Arabidopsis thaliana* Col-0 seeds were surface sterilized, stratified in the dark at 4°C for 4 days, and grown hydroponically to maturity (5 weeks) in Microclima environment-controlled cabinets (Snijders Scientific), maintaining 20°C, 55% relative humidity, and 12 h light (150 μE m^−2^ s^1^):12 h dark as described previously [72, 73]. 10-13 Arabidopsis rosettes were harvested and pooled at each sampling time. Harvesting occurred every 3 h over the last 24 h at 20°C, and on days 1 and 4 after transfer to 4°C giving 26 time-points in the time-series (Fig. 1a). Day 1 at 4°C represents the “transition” from 20°C to 4°C when plants first begin to experience the temperature decrease; Day 4 at 4°C represents “acclimation” where plants have been exposed to 4°C for 4 days (Fig. 1a). Three biological replicates were generated for each time-point in separate experiments (78 samples in total). The same growth cabinet was used for all repeats to eliminate the potential effects of minor changes in light intensities and light quality on gene expression. Additionally, to avoid interference in the experiment from possible microclimates within the growth cabinet, 26 trays for each time-point were placed in a randomised fashion. The switch to 4°C from 20°C was initiated at dusk. In a temperature reduction, the cabinet used here typically takes 2 h to reach 4°C air temperature (Additional file 1: Figure S6). Tissue was rapidly frozen in liquid N_2_ and stored at −80°C until isolation of RNA and preparation of cDNA.

### RNA Extraction

Total RNA was extracted from Arabidopsis tissue using RNeasy Plant Mini kit (Qiagen), followed by either on-column DNase treatment (for HR RT-PCR, see below), or the TURBO DNA-free™ kit (Ambion) (for library preparation and qRT-PCR, see below).

### Library preparation and sequencing

RNA-seq libraries were constructed for 78 RNA samples by following instructions for a TruSeq RNA library preparation (Illumina protocol 15026495 Rev. B). In these preparations, polyA selection was used to enrich for mRNA, RNA was fragmented for 8 min at 94°C, and random hexamers were used for first strand cDNA synthesis. The 78 libraries had an average insert size of approximately 280bp and each library was sequenced on three different sequencing lanes (27 lanes were used in total) of Illumina HiSeq 2500 platform generating 100 bp paired-end reads. The total number of raw reads generated in the RNA-seq data was 9.52 Bn paired-end reads giving approximately 360 M paired-end reads per time-point (Additional file 17: Table S15).

Residual adaptor sequences at both 5’ and 3’ ends were removed from raw reads using cutadapt version 1.4.2 (https://pypi.python.org/pypi/cutadapt/1.4.2) with quality score threshold set at 20 and minimum length of the trimmed read kept at 20. The “--paired-output” option was turned on to keep the two paired read files synchronized and avoid unpaired reads. The sequencing files before and after the trimming were examined using fastQC version 0.10.0.

### Quantification of transcripts and AS

Arabidopsis transcript expression from our RNA-seq experiment was carried out using Salmon version 0.82 [26] in conjunction with AtRTD2-QUASI augmented by 8 genes that were not originally present [27]. For indexing, we used the quasi mapping mode to build an auxiliary k-mer hash over k-mers of length 31 (--type quasi –k 31). For quantification, the option to correct for the sequence specific bias (“--seqBias”) was turned on. The number of bootstraps was set to 30 and all other parameters were on default settings. AtRTD2-QUASI is a modification of AtRTD2, a high quality reference transcript dataset for Arabidopsis thaliana Col-0 containing >82k unique transcripts, designed for the quantification of transcript expression [27]. Their use in validation of transcript structures and accurate quantification of individual transcript abundances for alternative splicing analyses was demonstrated previously using the biological repeats of two of the time-points analysed here [27]. Transcript expression results are in Additional file 18: Table S16.

### Differential gene expression (DE) and AS (DAS) analysis of the RNA-seq data

To carry out differential expression analysis, transcript quantification results generated by Salmon were processed and refined in successive steps (Additional file 1: Figure S18). First, transcript and gene read counts were generated from TPM data correcting for possible gene length variations across samples using tximport version 0.99.2 R package with the option “lengthScaledTPM” [74]. Second, read count data from sequencing replicates were summed for each biological sample. Third, genes and transcripts that are expressed at very low levels were removed from downstream analysis. The definition of a low expressed gene and transcript was determined by analysing mean-variance relationships [30]. The expected decreasing trend between the means and variances was observed in our data when removing transcripts that did not have ≥ 1 counts per million (CPM) in 3 or more samples out of 78, which provided an optimal filtering for low expression transcripts. At the gene level, if any transcript passed the expression level filtering step, the gene was included as an expressed gene and then the normalisation factor, which accounted for the raw library size, was estimated using the weighted trimmed Mean of M values method using edgeR version 3.12.1 [75]. Principal Component Analysis showed significant batch effects within the three biological replicates. Thus, batch effects between biological repeats were estimated using RUVSeq R package version 1.4.0 with the residual RUVr approach [76]. Normalised read counts in CPM were then log_2_ transformed and mean-variance trends were estimated and weights of variance adjustments were generated using the voom function in limma version 3.26.9 [30-32].

General linear models to determine differential expression at both gene and transcript levels were established and 18 contrast groups were set up where corresponding time-points in the day 1 and day 4 at 4°C blocks were compared to those of the 20°C block (e.g. block2.T1 vs block1.T1, block2.T2 vs block1.T2 etc; Additional file 1: Figures S1 and S2). P-values were adjusted for multiple testing [77]. For differential alternative splicing (DAS) analysis, the log_2_-fold changes (L_2_FCs) of each individual transcript of the gene were compared to the gene level L_2_FC, which is the average of all the transcripts weighted on standard deviation of each transcript. The consistency of expression changes among all the transcripts and the changes of expression at the gene level were tested using an overall F-test for the same 18 contrasts using the DiffSplice function (Additional file 1: Figure S2) [32]. For differential transcript usage (DTU) analysis, L_2_FCs of each transcript from DAS genes were compared to the L_2_FC of the weighted average of all the other transcripts from the same gene. The consistency of expression changes between each transcript and the change of expression of the other transcripts were tested using a t-test for the same 18 contrasts using the DiffSplice function (Additional file 1: Figure S2). DTU transcripts which coded for protein isoforms were determined from transcript translations of AtRTD2 [27].

Genes were significantly DE at the gene level if they had at least two contrast groups at consecutive time-points with adjusted *p*<0.01 and ≥2-fold change in expression in each contrast group (Additional file 1: Figure S1). Genes/transcripts with significant DAS/DTU, had at least two consecutive contrast groups with adjusted *p*<0.01 and with these contrast groups having at least one transcript with ≥10% change in expression (Additional file 1: Figure S2). Gene functional annotation was performed using R package RDAVIDWebService version 1.8.0 [78-80]. The possibility of a gene and transcript being identified by accident as cold responsive by the statistical method was tested. Time-points T1 and T9 (Fig. 1a) are virtually identical as they both represent dusk samples at 20°C, the only difference being they are 24 hours apart, such that few DE or DAS genes and DTU transcripts were expected when comparing these time-points. Indeed, no significant DE or DAS gene, nor DTU transcript, was identified between T1 and T9. This suggests our statistical method to select cold-responsive genes and transcripts is conservative and controls the number of false positives.

### Identification of isoform switches

5,317 high abundance transcripts, whose average expression accounts for >20% of total gene expression at at least one time-point, were selected from DAS gene transcripts for the isoform switch analysis using the TSIS R package which is a tool to detect significant transcript isoform switches in time-series data [45]. Switches between any two time-points were identified by using the default parameters in which i) the probability of switch (i.e. the frequency of samples reversing their relative abundance at the switches) was set to >0.5; ii) the sum of the average differences of the two isoforms in both intervals before and after the switch point were set at ΔTPM>1; iii) the significance of the differences between the switched isoform abundances before and after the switch was set to *p*<0.001; and iv) both intervals before and after switch must consist of at least 2 consecutive time-points in order to detect long lasting switches. SUPPA version 2.1 [46] was then used to identify the specific AS events (e.g. intron retention, alternative 3’ or 5’ splice site selection, exon skip) that distinguished the pair of switch transcript isoforms.

### Quantitative reverse transcription RT-PCR (qRT-PCR)

Real Time RT-PCR was performed essentially as described previously [72, 73]. Complementary DNA (cDNA) was synthesised from 2 μg of total RNA using oligo dT primers and SuperScriptII reverse transcriptase (ThermoFisher Scientific). Each reaction (1:100 dilution of cDNA) was performed with Brilliant III SYBR Green QPCR Master Mix (Agilent) on a StepOnePlus (Fisher Scientific-UK Ltd, Loughborough, UK) real-time PCR system. The average Ct values for PP2A (AT1G13320) and IPP2 (AT3G02780) were used as internal control expression levels. The delta-delta Ct algorithm [81] was used to determine relative changes in gene expression. Primer sequences are provided in Supporting Information Table 17a.

### High-resolution (HR) RT-PCR

HR RT-PCR reactions were conducted as described previously [82]. Gene-specific primer pairs were used for analysing the expression and alternative splicing of different genes (Additional file 19: Table S17). For each primer pair, the forward primer was labelled with 6-carboxyfluorescein (FAM). cDNA was synthesised from 4 μg of total RNA using the Sprint RT Complete – Double PrePrimed kit following manufacturer’s instructions (Clontech Laboratories, Takara Bio Company, USA). The PCR reaction usually contained 3 μL of diluted cDNA (1:10) as a template, 0.1 μL of each of the forward and reverse primers (100 mM), 2 μL of 10 X PCR Buffer, 0.2 μL of Taq Polymerase (5U/μL, Roche), 1 μL of 10 mM dNTPs (Invitrogen, Life Technologies) and RNase-free water (Qiagen) up to a final volume of 20 μL. For each reaction, an initial step at 94°C for 2 min was used followed by 24-26 cycles of 1) denaturation at 94°C for 15 sec, 2) annealing at 50°C for 30 sec and 3) elongation at 70°C for either 1 min (for fragments smaller than 1000 bp) or 1.5 min (for fragments between 1000-1200 bp) and a final extension cycle of 10 min at 70°C.

To separate the RT-PCR products, 1.5 **u**L of PCR product was mixed with 8.5 μL of Hi-Di^TM^ formamide (Applied Biosystems) and 0.01 μL of GeneScan^TM^ 500 LIZ^TM^ dye or 0.04 μL of GeneScanTM 1200 LIZ^TM^ dye size standard and run on a 48-capillary ABI 3730 DNA Analyser (Applied Biosystems, Life Technologies). PCR products were separated to single base-pair resolution and the intensity of fluorescence was measured and used for quantification in Relative Fluorescent Units (RFU). The different PCR products and their peak levels of expression were calculated using the Genemapper^®^ software (Applied Biosystems, Life Technologies).

### Identification and characterisation of the *u2b”-like* mutant

cDNA was synthesised as described above for HR RT-PCR. PCR was performed using cDNAs and GoTaq Green DNA polymerase (Promega) following manufacturer’s instructions. Primer sequences are provided in Additional file 19: Table S17.

### Freezing and electrolyte leakage assay

Cold acclimated plants were assessed for damage after freezing conditions. Sterilised seeds were sown on MS-agar plates and after 7 days seedlings were transferred to peat plugs for growth in 12:12 light-dark cycles, 150 to 200 μE/m^2^/s at 20°C for 4 weeks. Plants were then transferred to 5°C, 10:14 LD cycles (150 μE/m^2^/s) for ca. 14 d after which they were used in either a qualitative or quantitative assay. In the qualitative assay, cold-acclimated plants were transferred at dusk to either −8.0°C or −8.5°C for 24 h, then transferred to 5°C, 10:14 LD cycles (150 μE/m^2^/s) for 24 h and finally to 12:12 light-dark cycles, 150 to 200 μE/m^2^/s at 20°C for 1 week after which they were assessed for signs of regrowth, indicating survival. In the quantitative assay, we performed the electrolyte leakage test [83]. In brief, three leaf discs were collected from each cold-acclimated plant, forming a pseudo-replicate. For each temperature and genotype, six pseudo-replicates were obtained, representing one biological replicate (Additional file 1: Figure S19). Ice nucleation was initiated in individual test tubes for each pseudo-replicate by introducing ice chips and tubes were cooled progressively to the sub-zero temperatures indicated. Conductivity measurements were made after thawing and then again after complete loss of all electrolytes, to give a percentage measurement of electrolyte loss in each sample. In total 4 biological replicates were analysed. Percentage ion leakage data were first divided by 100 and then square-root and arc-sine transformed before analysis in one-tailed t-tests.

## Acknowledgements

This work was supported by funding from the Biotechnology and Biological Sciences Research Council (BBSRC) [BB/K006568/1, BB/P009751/1, BB/N022807/1 to JWSB; BB/K006835/1 to HGN; BB/M010996/1 – EASTBIO Doctoral Training Partnership for JCE]; the Scottish Government Rural and Environment Science and Analytical Services division (RESAS) [to JWSB and RZ].

We acknowledge the European Alternative Splicing Network of Excellence (EURASNET) [LSHG-CT-2005-518238] for catalysing important collaborations. We thank Janet Laird (University of Glasgow) for technical assistance, Katherine Denby and Iulia Gherman (University of York) for the list of Arabidopsis transcription factors. RNA-sequencing was performed at The Genome Analysis centre (now Earlham Institute), Norwich. We thank Robbie Waugh and Piers Hemsley for critical reading of the manuscript and helpful comments. We wish to apologize to all the authors whose relevant work was not cited in this article due to space limitations.

## Additional files

**File name:** Additional file 1

**File format:** .pdf

**Title of data:** Supplementary figures

**Description of data:**

**Figure S1.** Differential expression (DE) analysis.

**Figure S2.** Differential alternative splicing (DAS) and transcript usage (DTU) analysis.

**Figure S3.** Number of genes that are DE and DAS at each time-point in Day 1 and Day 4 at 4°C when compared to the 20°C control.

**Figure S4.** Comparison of DE and DAS genes between Day 1 and Day 4 at 4°C.

**Figure S5.** Expression profiles of CBFs, selected COR genes and known cold response genes.

**Figure S6.** Graph of air temperature reduction over time inside Microclima growth cabinet (Snijders Scientific).

**Figure S7.** Hierarchical clustering and heat map of Arabidopsis DE cold responsive genes and key GO terms.

**Figure S8.** Average expression profiles of DE gene clusters from heatmap in Figure S7.

**Figure S9.** Expression profiles of cold response DE-only genes.

**Figure S10.** Average expression profiles of DTU transcript clusters from heatmap in Fig. 3.

**Figure S11.** Expression profiles of cold response DE+DAS genes.

**Figure S12.** Expression profiles of cold response DAS-only genes.

**Figure S13.** Sensitivity of AS to small reductions in temperature.

**Figure S14.** Rapid changes in AS in response to gradual decrease in temperature from 20°C to 4°C in the first 0-3 hours of the cold treatment.

**Figure S15.** Identification and characterisation of *u2b’’-like* mutant.

**Figure S16.** Alignment of protein sequences of U2B”-LIKE and U2B’’ genes in six Brassicaceae species.

**Figure S17.** AS events significantly affected in the *u2b”-like* knock-out plants compared to Col-0 (WT).

**Figure S18.** RNA-seq data pre-processing pipeline.

**Figure S19.** Electrolyte leakage assay experimental design.

**File name:** Additional file 2

**File format:** .xlsx

**Title of data:** Table S1

**Description of data:** Table S1a. Number of genes and transcripts from results of the analysis of differentially expressed, differentially alternatively spliced and differential transcript usage. Table S1b. Expressed genes and transcripts in RNA-seq time course, as well as genes differentially regulated by cold at the expression (DE) and/or alternative splicing level (DAS). Table S1c. DE and DAS genes identified at Day 1 and/or Day4.

**File name:** Additional file 3

**File format:** .xlsx

**Title of data:** Table S2

**Description of data:** Gene lists of the up- and down-regulated DE genes in the different contrast groups.

**File name:** Additional file 4

**File format:** .pdf

**Title of data:** Supplementary Table S3

**Description of data:** Summary of previous work on wild-type Arabidopsis genome-wide cold response.

**File name:** Additional file 5

**File format:** .xlsx

**Title of data:** Table S4

**Description of data:** Gene lists of known Arabidopsis cold-response genes in Table S5.

**File name:** Additional file 6

**File format:** .xlsx

**Title of data:** Table S5

**Description of data:** Novel genes differentially regulated by cold at the expression (DE) and/or alternative splicing level (DAS).

**File name:** Additional file 7

**File format:** .pdf

**Title of data:** Supplementary Notes

**Description of data:** Additional information is provided on 1) Comparison of cold response experimental conditions. 2) U2B” and U2B”-LIKE genes in animals and plants. 3) Reference gene lists of RNA-binding, splicing factor and spliceosomal protein and transcription factor genes.

**File name:** Additional file 8

**File format:** .xlsx

**Title of data:** Table S6

**Description of data:** Table S6a. Gene Ontology (GO) enrichment analysis of DE genes (showing only significant terms with FDR < 0.05). Table S6b. Gene Ontology (GO) enrichment analysis of DAS genes (showing only significant terms with FDR < 0.05).

**File name:** Additional file 9

**File format:** .xlsx

**Title of data:** Table S7

**Description of data:** Table S7a. Genes in DE heatmap clusters. Table S7b. DTU transcripts in heatmap clusters. Table S7c. Gene Ontology (GO) enrichment analysis of individual heatmap DE gene clusters (showing only significant terms with FDR<0.05) (see Additional file 1: Figure S7).

**File name:** Additional file 10

**File format:** .xlsx

**Title of data:** Table S8

**Description of data:** Table S8a. Gene lists of 7,302 DE genes organised by the time-point at which they first become significantly differentially expressed upon cold. Table S8b. Gene descriptions of the 648 genes which are first detected as significantly differentially expressed at 3 hours after the start of cold treatment (T10) in Table S8a. Table S8c. Gene descriptions of the 1452 genes which are first detected as significantly differentially expressed at 6 hours after the start of cold treatment (T11) in Table S8a. Table S8d. Gene lists of 2,442 DAS genes organised by the time-point at which they first become significantly differentially alternatively spliced upon cold. Table S8e. Gene descriptions of the 388 genes which are first detected as significantly differentially alternatively spliced at 3 hours after the start of cold treatment (T10) in Table S8d. Table S8f. Gene descriptions of the 768 genes which are first detected as significantly differentially alternatively spliced at 6 hours after the start of cold treatment (T11) in Table S8d.

**File name:** Additional file 11

**File format:** xlsx

**Title of data:** Table S9

**Description of data:** Table S9a. DAS gene lists organised by different ΔPS cut-off values. Table S9b. Gene descriptions of 1110 DAS genes with ΔPS >0.2 in Table S9a. Table S9c. Gene descriptions of 390 DAS genes with ΔPS >0.3 in Table S9a.

**File name:** Additional file 12

**File format:** .xlsx

**Title of data:** Table S10

**Description of data:** Table S10a. Isoform switch analysis of transcripts from DAS genes using the TSIS package (Guo et al., 2017). Table S10b. AS events identified with SUPPA (Alamancos et al., 2015) for each isoform switch. Table S10c. Gene lists of DAS genes with Isoform Switches (IS) between 0-3 h cold (T9-T10), 3-6 h cold (T10-T11) and 0-6 h cold (T9-T11).

**File name:** Additional file 13

**File format:** .xlsx

**Title of data:** Table S11

**Description of data:** Table S11a. List of 2534 Arabidopsis Transcription Factor genes. Table S11b. List of 798 Arabidopsis SF-RBP genes.

**File name:** Additional file 14

**File format:** .xlsx

**Title of data:** Table S12

**Description of data:** Table S12a. Summary of 271 TF genes differentially regulated by cold at the alternative splicing level (DAS) and TF genes with the earliest and largest changes in AS. Table S12b. Summary of 166 SF-RBP genes differentially regulated by cold at the alternative splicing level (DAS) and SF-RBP genes with the earliest and largest changes in AS. Table S12c. TF and SF-RBP DAS genes with the largest and most rapid changes in AS. Table S12d. TF and SF-RBP DE genes with the largest and most rapid changes in expression. Table S12e. Summary of functions/classifications of TF genes showing the largest and most rapid changes in gene expression and/or AS in cold treatment. Table S12f. Classifications of SF-RBPs showing the largest and most rapid changes in gene expression and/or AS in cold treatment.

**File name:** Additional file 15

**File format:** .xlsx

**Title of data:** Table S13

**Description of data:** Table S13a. Timing of significantly differentially expressed (DE) SF-RBP genes in the first 12 h after onset of cold treatment (T9-T13). Table S13b. Timing of significantly differentially expressed (DE) TF genes in the first 12 h after onset of cold treatment (T9-T13). Table S13c. Timing of significantly differentially alternatively spliced (DAS) SF-RBP genes in the first 12 h after the onset of cold treatment (T10-T13). Table S13d. Timing of significantly differentially alternatively spliced (DAS) TF genes in the first 12 h after the onset of cold treatment (T10-T13).

**File name:** Additional file 16

**File format:** .xlsx

**Title of data:** Table S14

**Description of data:** AS data for the *u2b”-like* HR RT-PCR analysis panel.

**File name:** Additional file 17

**File format:** .xlsx

**Title of data:** Table S15

**Description of data:** Summary of RNA-seq data derived from Arabidopsis rosettes time-series cold response study.

**File name:** Additional file 18

**File format:** .xlsx

**Title of data:** Table S16

**Description of data:** Transcript expression in the Arabidopsis cold response.

**File name:** Additional file 19

**File format:** .xlsx

**Title of data:** Table S17

**Description of data:** Table S17a. Primers used for the speed and sensitivity of AS using Real Time RT-PCR. Table S17b. Primers used for the speed and sensitivity of AS using HR RT-PCR. Table S17c. Primers used for the HR RT-PCR analysis in the u2b’’-like mutant. Table S17d. Primers used for the amplification of the U2B (AT2G30260) and U2B’’-LIKE (AT1G06960) genes in the SALK_060577 line using RT-PCR.

## References

1. Thomashow MF: Molecular basis of plant cold acclimation: insights gained from studying the CBF cold response pathway. Plant Physiology 2010, 154:571–577.

2. Zhu JK: Abiotic Stress Signaling and Responses in Plants. Cell 2016, 167:313–324.

3. Knight MR, Knight H: Low-temperature perception leading to gene expression and cold tolerance in higher plants. New Phytologist 2012, 195:737–751.

4. Kim JM, Sasaki T, Ueda M, Sako K, Seki M: Chromatin changes in response to drought, salinity, heat, and cold stresses in plants. Frontiers in Plant Science 2015, 6:114.

5. Barrero-Gil J, Salinas J: Post-translational regulation of cold acclimation response. Plant Science 2013, 205-206:48–54.

6. Jia Y, Ding Y, Shi Y, Zhang X, Gong Z, Yang S: The cbfs triple mutants reveal the essential functions of CBFs in cold acclimation and allow the definition of CBF regulons in Arabidopsis. New Phytologist 2016, 212:345–353.

7. Zhao C, Zhang Z, Xie X, Si T, Li Y, Zhu JK: Mutational Evidence for the Critical Role of CBF Transcription Factors in Cold Acclimation in Arabidopsis. Plant Physiology 2016, 171:2744–2759.

8. Park S, Lee CM, Doherty CJ, Gilmour SJ, Kim Y, Thomashow MF: Regulation of the Arabidopsis CBF regulon by a complex low-temperature regulatory network. Plant Journal 2015, 82:193–207.

9. Harmer SL: The circadian system in higher plants. Annual Review of Plant Biology 2009, 60:357–377.

10. Laloum T, Martín G, Duque P: Alternative Splicing Control of Abiotic Stress Responses. Trends in Plant Science 2017, [Epub ahead of print].

11. Staiger D, Brown JW: Alternative splicing at the intersection of biological timing, development, and stress responses. Plant Cell 2013, 25:3640–3656.

12. Leviatan N, Alkan N, Leshkowitz D, Fluhr R: Genome-wide survey of cold stress regulated alternative splicing in Arabidopsis thaliana with tiling microarray. PLoS One 2013, 8:e66511.

13. Li S, Yamada M, Han X, Ohler U, Benfey PN: High-Resolution Expression Map of the Arabidopsis Root Reveals Alternative Splicing and lincRNA Regulation. Developmental Cell 2016, 39:508–522.

14. Pajoro A, Severing E, Angenent GC, Immink RGH: Histone H3 lysine 36 methylation affects temperature-induced alternative splicing and flowering in plants. Genome Biology 2017, 18:102.

15. Hartmann L, Drewe-Boß P, Wießner T, Wagner G, Geue S, Lee HC, Obermüller DM, Kahles A, Behr J, Sinz FH, et al: Alternative Splicing Substantially Diversifies the Transcriptome during Early Photomorphogenesis and Correlates with the Energy Availability in Arabidopsis. Plant Cell 2016, 28:2715–2734.

16. Klepikova AV, Kasianov AS, Gerasimov ES, Logacheva MD, Penin AA: A high resolution map of the Arabidopsis thaliana developmental transcriptome based on RNA-seq profiling. Plant Journal 2016, 88:1058–1070.

17. Mastrangelo AM, Marone D, Laidò G, De Leonardis AM, De Vita P: Alternative splicing: enhancing ability to cope with stress via transcriptome plasticity. Plant Science 2012, 185-186:40–49.

18. Lee Y, Rio DC: Mechanisms and Regulation of Alternative Pre-mRNA Splicing. Annual Review of biochemistry 2015, 84:291–323.

19. Fu XD, Ares MJ: Context-dependent control of alternative splicing by RNA-binding proteins. Nature Reviews Genetics 2014, 15:689–701.

20. Marquez Y, Brown JW, Simpson C, Barta A, Kalyna M: Transcriptome survey reveals increased complexity of the alternative splicing landscape in Arabidopsis. Genome Research 2012, 22:1184–1195.

21. Zhang X-N, Mount SM: Two alternatively spliced isoforms of the Arabidopsis SR45 protein have distinct roles during normal plant development. Plant Physiology 2009, 150:1450–1458.

22. Schlaen RG, Mancini E, Sanchez SE, Perez-Santángelo S, Rugnone ML, Simpson CG, Brown JW, Zhang X, Chernomoretz A, Yanovsky MJ: The spliceosome assembly factor GEMIN2 attenuates the effects of temperature on alternative splicing and circadian rhythms. Proceedings of the National Academy of Sciences of the United States of America 2015, doi/10.1073/pnas.1504541112.

23. Cheng C, Wang Z, Yuan B, Li X: RBM25 Mediates Abiotic Responses in Plants. Frontiers in Plant Science 2017, 8:292.

24. Reddy AS, Marquez Y, Kalyna M, Barta A: Complexity of the alternative splicing landscape in plants. Plant Cell 2013, 25:3657–3683.

25. Bray N, Pimentel H, Melsted P, Pachter L: Near-optimal probabilistic RNA-seq quantification. Nature Biotechnology 2016, 34:525–527.

26. Patro R, Duggal G, Love MI, Irizarry RA, Kingsford C: Salmon provides fast and bias-aware quantification of transcript expression. Nature Methods 2017, 14:417–419.

27. Zhang R, Calixto CPG, Marquez Y, Venhuizen P, Tzioutziou NA, Guo W, Spensley M, Entizne JC, Frei dit Frey N, Hirt H, et al: A high quality Arabidopsis transcriptome for accurate transcript-level analysis of alternative splicing. Nucleic Acids Research 2017, 45:5061–5073.

28. Filichkin SA, Priest HD, Givan SA, Shen R, Bryant DW, Fox SE, Wong WK, Mockler TC: Genome-wide mapping of alternative splicing in Arabidopsis thaliana. Genome Res 2010, 20:45–58.

29. Ding F, Cui P, Wang Z, Zhang S, Ali S, Xiong L: Genome-wide analysis of alternative splicing of pre-mRNA under salt stress in Arabidopsis. BMC Genomics 2014, 15:431.

30. Law CW, Alhamdoosh M, Su S, Smyth GK, Ritchie ME: RNA-seq analysis is easy as 1-2-3 with limma, Glimma and edgeR. F1000 Research 2016, 5.

31. Law CW, Chen Y, Shi W, Smyth GK: voom: Precision weights unlock linear model analysis tools for RNA-seq read counts. Genome Biology 2014, 15:R29.

32. Ritchie ME, Phipson B, Wu D, Hu Y, Law CW, Shi W, Smyth GK: limma powers differential expression analyses for RNA-sequencing and microarray studies. Nucleic Acids Research 2015, 43:e47.

33. Vogel JT, Zarka DG, Van Buskirk HA, Thomashow MF: Roles of the CBF2 and ZAT12 transcription factors in configuring the low temperature transcriptome of Arabidopsis. Plant Journal 2005, 41:195–211.

34. Carvallo MA, Pino MT, Jeknic Z, Zou C, Doherty CJ, Shiu SH, Chen TH, Thomashow MF: A comparison of the low temperature transcriptomes and CBF regulons of three plant species that differ in freezing tolerance: Solanum commersonii, Solanum tuberosum, and Arabidopsis thaliana. Journal of Experimental Botany 2011, 62:3807–3819.

35. Gehan MA, Park S, Gilmour SJ, An C, Lee CM, Thomashow MF: Natural variation in the C-repeat binding factor cold response pathway correlates with local adaptation of Arabidopsis ecotypes. Plant Journal 2015, 84:682–693.

36. Gilmour SJ, Zarka DG, Stockinger EJ, Salazar MP, Houghton JM, Thomashow MF: Low temperature regulation of the Arabidopsis CBF family of AP2 transcriptional activators as an early step in cold-induced COR gene expression. Plant Journal 1998, 16:433–442.

37. Guan Q, Wu J, Zhang Y, Jiang C, Liu R, Chai C, Zhu J: A DEAD box RNA helicase is critical for pre-mRNA splicing, cold-responsive gene regulation, and cold tolerance in Arabidopsis. Plant Cell 2013, 25:342–356.

38. Kidokoro S, Maruyama K, Nakashima K, Imura Y, Narusaka Y, Shinwari ZK, Osakabe Y, Fujita Y, Mizoi J, Shinozaki K, Yamaguchi-Shinozaki K: The phytochrome-interacting factor PIF7 negatively regulates DREB1 expression under circadian control in Arabidopsis. Plant Physiology 2009, 151:2046–2057.

39. Wang F, Guo Z, Li H, Wang M, Onac E, Zhou J, Xia X, Shi K, Yu J, Zhou Y: Phytochrome A and B Function Antagonistically to Regulate Cold Tolerance via Abscisic Acid-Dependent Jasmonate Signaling. Plant Physiology 2016, 170:459–471.

40. Lee CM, Thomashow MF: Photoperiodic regulation of the C-repeat binding factor (CBF) cold acclimation pathway and freezing tolerance in Arabidopsis thaliana. Proceedings of the National Academy of Sciences of the United States of America 2012, 109:15054–15059.

41. Krall L, Reed JW: The histidine kinase-related domain participates in phytochrome B function but is dispensable. Proceedings of the National Academy of Sciences of the United States of America 2000, 97:8169–8174.

42. Müller R, Fernández AP, Hiltbrunner A, Schäfer E, Kretsch T: The histidine kinase-related domain of Arabidopsis phytochrome a controls the spectral sensitivity and the subcellular distribution of the photoreceptor. Plant Physiology 2009, 150:1297–1309.

43. Ding L, Kim SY, Michaels SD: FLOWERING LOCUS C EXPRESSOR family proteins regulate FLOWERING LOCUS C expression in both winter-annual and rapid-cycling Arabidopsis. Plant Physiology 2013, 163:243–252.

44. Kim SY, Michaels SD: SUPPRESSOR OF FRI 4 encodes a nuclear-localized protein that is required for delayed flowering in winter-annual Arabidopsis. Development 2006, 133:4699–4707.

45. Guo W, Calixto CPG, Brown JWS, Zhang R: TSIS: an R package to infer alternative splicing isoform switches for time-series data. Bioinformatics 2017, 33:3308–3310.

46. Alamancos GP, Pagès A, Trincado JL, Bellora N, Eyras E: Leveraging transcript quantification for fast computation of alternative splicing profiles. RNA 2015, 21:1521–1531.

47. Rosembert M: The role of pre-mRNA splicing and splicing related proteins in the cold acclimation induced adjustment of photosynthesis and the acquisition of freezing tolerance in Arabiodpsis thaliana. University of Ottawa, Faculty of Science; 2017.

48. Hiller M, Zhang Z, Backofen R, Stamm S: Pre-mRNA secondary structures influence exon recognition. PLoS Genetics 2007, 3:e204.

49. Luco RF, Allo M, Schor IE, Kornblihtt AR, Misteli T: Epigenetics in alternative pre-mRNA splicing. Cell 2011, 144:16–26.

50. Berry S, Dean C: Environmental perception and epigenetic memory: mechanistic insight through FLC. Plant Journal 2015, 83:133–148.

51. Kim JM, To TK, Ishida J, Matsui A, Kimura H, Seki M: Transition of chromatin status during the process of recovery from drought stress in Arabidopsis thaliana. Plant and Cell Physiology 2012, 53:847–856.

52. Haak DC, Fukao T, Grene R, Hua Z, Ivanov R, Perrella G, Li S: Multilevel Regulation of Abiotic Stress Responses in Plants. Frontiers in Plant Science 2017, 8:1564.

53. Verhage L, Severing EI, Bucher J, Lammers M, Busscher-Lange J, Bonnema G, Rodenburg N, Proveniers MC, Angenent GC, Immink RG: Splicing-related genes are alternatively spliced upon changes in ambient temperatures in plants. PLoS One 2017, 12:e0172950.

54. Doherty CJ, Van Buskirk HA, Myers SJ, Thomashow MF: Roles for Arabidopsis CAMTA transcription factors in cold-regulated gene expression and freezing tolerance. Plant Cell 2009, 21:972–984.

55. Kanno T, Lin WD, Fu JL, Chang CL, Matzke AJM, Matzke M: A Genetic Screen for Pre-mRNA Splicing Mutants of Arabidopsis thaliana Identifies Putative U1 snRNP Components RBM25 and PRP39a. Genetics 2017, 207:1347–1359.

56. Barta A, Marquez Y, Brown JWS: Challenges in Plant Alternative Splicing. In Alternative pre-mRNA Splicing: Theory and Protocols. Edited by Stamm S, Smith CWJ, Lührmann R. Weinheim, Germany: Wiley-VCH Verlag GmbH & Co.; 2012

57. Bove J, Kim CY, Gibson CA, Assmann SM: Characterization of wound-responsive RNA-binding proteins and their splice variants in Arabidopsis. Plant Molecular Biology 2008, 67:71–88.

58. Savaldi-Goldstein S, Aviv D, Davydov O, Fluhr R: Alternative splicing modulation by a LAMMER kinase impinges on developmental and transcriptome expression. Plant Cell 2003, 15:926–938.

59. Medina MW, Gao F, Naidoo D, Rudel LL, Temel RE, McDaniel AL, Marshall SM, Krauss RM: Coordinately regulated alternative splicing of genes involved in cholesterol biosynthesis and uptake. PLoS One 2011, 6:e19420.

60. Mauger O, Lemoine F, Scheiffele P: Targeted Intron Retention and Excision for Rapid Gene Regulation in Response to Neuronal Activity. Neuron 2016, 92:1266–1278.

61. Preußner M, Goldammer G, Neumann A, Haltenhof T, Rautenstrauch P, Müller-McNicoll M, Heyd F: Body Temperature Cycles Control Rhythmic Alternative Splicing in Mammals. Molecular Cell 2017, 67:433–446.

62. Stamm S: Regulation of alternative splicing by reversible protein phosphorylation. The Journal of Biological Chemistry 2008, 283:1223–1227.

63. Jung JH, Domijan M, Klose C, Biswas S, Ezer D, Gao M, Khattak AK, Box MS, Charoensawan V, Cortijo S, et al: Phytochromes function as thermosensors in Arabidopsis. Science 2016, 354:886–889.

64. Jangi M, Sharp PA: Building robust transcriptomes with master splicing factors. Cell 2014, 159:487–498.

65. Fiszbein A, Kornblihtt AR: Alternative splicing switches: Important players in cell differentiation. Bioessays 2017, 39:1600157.

66. Teige M, Scheikl E, Eulgem T, Dóczi R, Ichimura K, Shinozaki K, Dangl JL, Hirt H: The MKK2 pathway mediates cold and salt stress signaling in Arabidopsis. Molecular Cell 2004, 15:141–152.

67. Zhao C, Wang P, Si T, Hsu CC, Wang L, Zayed O, Yu Z, Dong J, Tao WA, Zhu JK: MAP Kinase Cascades Regulate the Cold Response by Modulating ICE1 Protein Stability. Developmental Cell 2017, 43:618–629.

68. Li H, Ding Y, Shi Y, Zhang X, Zhang S, Gong Z, Yang S: MPK3- and MPK6-Mediated ICE1 Phosphorylation Negatively Regulates ICE1 Stability and Freezing Tolerance in Arabidopsis. Developmental Cell 2017, 43:630–642.

69. Razanau A, Xie J: Emerging mechanisms and consequences of calcium regulation of alternative splicing in neurons and endocrine cells. Cell and Molecular Life Sciences 2013, 70:4527–4536.

70. Rausin G, Tillemans V, Stankovic N, Hanikenne M, Motte P: Dynamic nucleocytoplasmic shuttling of an Arabidopsis SR splicing factor: role of the RNA-binding domains. Plant Physiology 2010, 153:273–284.

71. de la Fuente van Bentem S, Anrather D, Roitinger E, Djamei A, Hufnagl T, Barta A, Csaszar E, Dohnal I, Lecourieux D, Hirt H: Phosphoproteomics reveals extensive in vivo phosphorylation of Arabidopsis proteins involved in RNA metabolism. Nucleic Acids Research 2006, 34:3267–3278.

72. James AB, Monreal JA, Nimmo GA, Kelly CL, Herzyk P, Jenkins GI, Nimmo HG: The circadian clock in Arabidopsis roots is a simplified slave version of the clock in shoots. Science 2008, 322:1832–1835.

73. James AB, Syed NH, Bordage S, Marshall J, Nimmo GA, Jenkins GI, Herzyk P, Brown JW, Nimmo HG: Alternative splicing mediates responses of the Arabidopsis circadian clock to temperature changes. Plant Cell 2012, 24:961–981.

74. Soneson C, Love MI, Robinson MD: Differential analyses for RNA-seq: transcript-level estimates improve gene-level inferences. F1000 Research 2015, 4:1521.

75. Robinson MD, McCarthy DJ, Smyth GK: edgeR: a Bioconductor package for differential expression analysis of digital gene expression data. Bioinformatics 2010, 26:139–140.

76. Risso D, Nqai J, Speed TP, Dudoit S: Normalization of RNA-seq data using factor analysis of control genes or samples. Nature Biotechnology 2014, 32:896–902.

77. Benjamini Y, Hochberg Y: Controlling the False Discovery Rate: A Practical and Powerful Approach to Multiple Testing. Journal of the Royal Statistical Society 1995, 57:289–300.

78. Huang DW, Sherman BT, Lempicki RA: Systematic and integrative analysis of large gene lists using DAVID Bioinformatics Resources. Nature Protocols 2009, 4:44–57.

79. Huang DW, Sherman BT, Lempicki RA: Bioinformatics enrichment tools: paths toward the comprehensive functional analysis of large gene lists. Nucleic Acids Research 2009, 37:1–13.

80. Fresno C, Fernández EA: RDAVIDWebService: a versatile R interface to DAVID. Bioinformatics 2013, 29:2810–2811.

81. Livak KJ, Schmittgen TD: Analysis of relative gene expression data using real-time quantitative PCR and the 2(-Delta Delta C(T)) Method. Methods 2001, 25:402–408.

82. Simpson CG, Fuller J, Maronova M, Kalyna M, Davidson D, McNicol J, Barta A, Brown JW: Monitoring changes in alternative precursor messenger RNA splicing in multiple gene transcripts. Plant Journal 2008, 53:1035–1048.

83. Hemsley PA, Hurst CH, Kaliyadasa E, Lamb R, Knight MR, De Cothi EA, Knight H: The Arabidopsis mediator complex subunits MED16, MED14, and MED2 regulate mediator and RNA polymerase II recruitment to CBF-responsive cold-regulated genes. Plant Cell 2014, 26:465–484.

